# Defining the cellular origin of seminoma by transcriptional and epigenetic mapping to the normal human germline

**DOI:** 10.1101/2024.04.20.590406

**Authors:** Keren Cheng, Yasunari Seita, Eoin C. Whelan, Ryo Yokomizo, Young Sun Hwang, Antonia Rotolo, Ian D. Krantz, Maninder Kaur, Jill P. Ginsberg, Priti Lal, Xunda Luo, Phillip M. Pierorazio, Rebecca L. Linn, Sandra Ryeom, Kotaro Sasaki

## Abstract

Aberrant male germline development can lead to the formation of seminoma, a testicular germ cell tumor. Seminomas are biologically similar to primordial germ cells (PGCs) and many bear an isochromosome 12p [i(12p)] with two additional copies of the short arm of chromosome 12. By mapping seminoma transcriptomes and open chromatin landscape onto a normal human male germline trajectory, we find that seminoma resembles premigratory/migratory primordial germ cells, but exhibit enhanced germline and pluripotency programs, and upregulation of genes involved in apoptosis, angiogenesis, and MAPK/ERK pathways. Using pluripotent stem cell-derived PGCs from Pallister Killian syndrome patients mosaic for i(12p) to model seminoma, we identify gene dosage effects that may contribute to transformation. As murine seminoma models do not exist, our analyses provide critical insights into genetic, cellular and signaling programs driving seminoma transformation, and the newly developed in vitro platform permits evaluation of additional signals required for seminoma tumorigenesis.

## INTRODUCTION

Testicular germ cell tumors (GCTs) represent the most common cancer in young men and the incidence of GCTs is increasing worldwide^1,2^. Although the survival rate is generally favorable after chemotherapy, a number of patients succumb to the disease due to relapse and chemo-resistance, and many survivors suffer from life-long infertility and chemo-toxic side effects^3,4^. The most common histologic subtype of GCT is the seminoma, which exhibits morphologic and biological similarities to primordial germ cells (PGCs) or gonocytes, the precursors for both oocytes and spermatozoa^5,6^. Although previous studies reveal that seminomas exhibit overall hypotetraploidy of the genome, up to 90% bear the pathognomonic marker chromosome, isochromosome 12p [i(12p)], which contains two additional copies of the short arm of chromosome 12 (12p) ^6–9^, suggesting potential gene dosage effects. Likewise, a subset of seminomas is characterized by activating mutations in *KIT*, which encodes a cell surface receptor tyrosine kinase that drives downstream signaling through the RAS/MAPK pathway, potentially contributing to their transformation^10–12^. However, the precise genetic basis and biological characteristics of seminoma remain poorly defined, thereby impeding the development of targeted therapeutic strategies against this form of cancer.

Despite recent efforts to understand early germ cell development at a single cell resolution^13–18^, a complete trajectory of human PGC development into haploid spermatids has not yet been defined, making it difficult to determine the exact developmental stage from which seminoma diverge. Progress is further impeded by the lack of relevant mouse models, the paucity of appropriate in vitro models, the scarcity of seminoma samples, heterogeneity and heavy lymphocytic infiltrates within tumors that can confound interpretation of bulk tumor analyses. To overcome these hurdles, we first established a comprehensive lineage trajectory map of the spectrum of human male germline development, from migrating PGCs in the 4 week post fertilization (wpf) embryo to haploid spermatids in 12 year old testis. Using this as a reference, we successfully mapped seminoma single cell transcriptomes to the normal developmental coordinates, identifying premigratory/migratory PGCs as the divergent node. Leveraging the power of single cell RNA-seq and ATAC-seq, we also determined the chromatin accessibility landscape of seminoma in comparison to developing germ cells, thereby identifying 1) potential pathways contributing to transformation, 2) a unique pattern of primate-specific transposable element (TE) activation that could potentially regulate gene programs driving human seminoma, and 3) a distinct molecular signature of the seminoma tumor microenvironment (TME). Finally, by generating induced pluripotent stem cell (iPSC)-derived PGCs from Pallister Killian syndrome patients mosaic for i(12p)^1^^9^, we have established an in vitro seminoma model to determine the impact of i(12p) on genome-wide gene expression.

## RESULTS

### A comprehensive single cell transcriptome atlas of human male germline development

Incomplete understanding of the lineage trajectory and gene expression dynamics that occurs during human male germline development, particularly within the fetal and prepubertal stages, currently restricts our ability to define the genetic and molecular underpinnings of seminoma. To overcome these limitations, we collected testes samples from the 2^nd^ and 3^rd^ trimester fetuses, an infant, as well as clinically prepubertal (5 and 8, and 12 year old) boys. Combined with previously collected data from 1^st^ and 2^nd^ trimester fetal samples^17,20^, this represents the most comprehensive human testicular sample set to date (Table S1). Single cell suspensions were subjected to single-cell RNA-seq (scRNA-seq) using a 10x Genomics platform. After filtering out low-quality cells, 133,725 cells remained for downstream analysis (Fig.S1A-B). Dimension reduction and projection of all transcriptomes into UMAP revealed 12 cell types, which were successfully annotated by differentially expressed genes (DEGs), including known markers of each cell type (Fig. S1C, D, Table S2). Among these, germ cells (cell type 1, 8363 cells) that were identified by a combination of specific marker genes (e.g., *NANOG*, *PDPN*, *DDX4*, *PRM1*) were parsed out for re-clustering analyses (Fig. S1E). Analyses of isolated germ cells projected onto the diffusion map yielded a comprehensive human male germline lineage trajectory, in which 18 distinct cell types identified by specific marker genes we and others previously identified using limited sample sets^17^ were aligned along a pseudo-time axis, that faithfully reflected developmental stages observed at the time points samples were collected (Fig. 1A-C). We identified M-prospermatogonia (M.pSpg) (cell type 3, *POU5F1*^+^*TFAP2C*^+^), Intermediate-prospermatogonia (Int.pSpg) (cell type 4, *ASB9*^+^) and T1-prospermatogonia (T1.pSpg) (cell type 5, *POU5F1*^−^*TFAP2C*^−^), which emerged at 6, 12, 15 wpf, respectively (Fig. 1D, Table S3)^17,18^. During the transition from M.pSpg to T1.pSpg through Int.pSpg, pluripotency-associated genes (e.g., *POU5F1*, *NANOG*, *TCL1A*) and germ cell specifier genes (e.g., *TFAP2C*, *PRDM1*, *SOX15*, *NANOS3*) were gradually turned off and the *MKI67* proliferation index decreased. In addition, a number of spermatogonial marker genes^14,15,17,21^, some encoding transcription factors (TFs), were upregulated (e.g., *SIX1*, *ESX1*, *EGR4*, *POU3F2*, *ID1*, *ID2*, *MAGEC2*, *CTAG2*, *PIWIL4*, *RHOXF1*) with DEGs enriched for GO terms such as “piRNA metabolic process” or “positive regulation of transcription from RNA polymerase II promoter” (Fig. 1D-F, S1F, S2A-I, Table S3). Notably, we identified two additional cell types that appeared earlier than M.pSpg in pseudotime, and were therefore annotated as early PGCs (PGCs.E) and late PGCs (PGCs.L) (Fig. 1A). These cell types were primary derived from 4-6 wpf (PGCs.E), or 8-9 wpf samples (PGCs.L) and thus represent PGCs at a migration/immediate early gonadal colonization stage and post-colonization stage, respectively (Table S4)^20^. Accordingly, PGCs.E upregulated genes related to “regulation of cell migration” (e.g., *NEXN*) and highly expressed pluripotency-associated and germ cell specifier genes, the expression of which showed modest downregulation as these cells transitioned into M.pSpg (Fig. 1D, S2A). PGCs.E also showed DEGs enriched with GO terms such as “aerobic respiration”, “oxidative phosphorylation”, consistent with previous reports of high oxidative phosphorylation activity in mouse PGCs (Table S3)^22^. During the PGCs.E to M.pSPg transition, genes related to spermatogenesis and meiosis (e.g., *TDRD9*, *TEX14*, *TEX15*), and cell cycle regulation (e.g., *MKI67*, *CDC25A*, *CDKN2C*) were upregulated (Fig. 1D). Notably, DNA methylation sensitive genes known as “germline genes” (e.g., *DDX4*, *DAZL*, *MAEL*)^23,24^ were also upregulated during this transition, which is consistent with ongoing DNA demethylation upon gonadal colonization (Fig. 1D-F, S2A)^25^.

**Figure 1.**
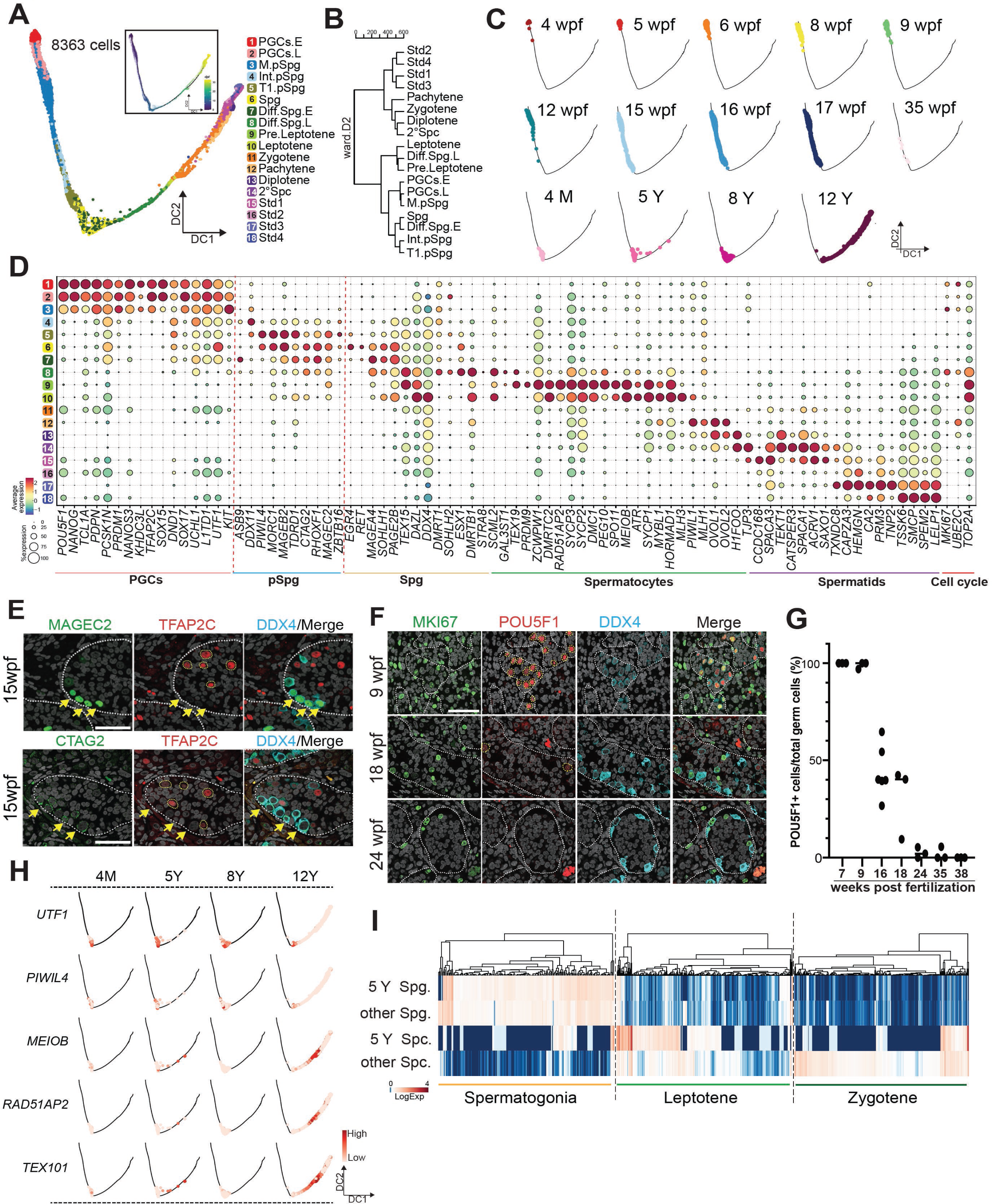
Single cell transcriptome of human male germline development. **(A)** DiffusionMap trajectory analysis of human male germline single cell transcriptomes covering from primordial germ cells (PGCs) to round spermatids (a total of 8363 cells). Based on DEGs and marker gene expression, cell clusters were annotated and assigned with unique colors. DiffusionMap trajectory colored by pseudotime (dpt) (inset). **(B)** Hierarchical Clustering of averaged expression (pseudo-bulk) values of annotated germ cell types using ward.D2. Top 2000 variably expressed genes were used. **(C)** DiffusionMap trajectory showing sample origins. Wpf, week post fertilization; M, months old; Y, years old. **(D)** Key marker gene expression selected from DEGs of each male germ cell type (related to Table S3). The dot size indicates the percentage of cells expressing the gene within each cell type. The color code indicates the relative average (pseudo-bulk) expression values. Gene expression is log normalized and then Z-score transformed. **(E)** Immunofluorescence (IF) images of testicular sections at 15 wpf for MAGEC2, CTAG2 (green), TFAP2C (red), DDX4 (cyan) merged with DAPI staining (white). Merged images of all four channels are shown on the right. Scale bar, 50 μm. **(F)** IF images of testicular sections at indicated stages for MKI67 (green), POU5F1 (Red), DDX4 (cyan) merged with DAPI (white). Merged images of all four channels are shown on the right. Scale car, 50 μm. **(G)** IF quantification showing the proportion of POU5F1^+^ PGCs/M.pSpg among all germ cells (DDX4^+^ or POU5F1^+^) in indicated fetal stages. At least 3-6 randomly selected images from 1-2 embryos per stage are counted. **(H)** Diffusion Map embeddings at indicated developmental stages. Expression of T1.pSpg/Spg markers (*UTF1*, *PIWIL4*) and Leptotene markers (*MEIOB*, *RAD51AP2*, *TEX101*) are shown. The color bar indicate Log normalized expression. **(I)** Heatmap showing expression of spermatogonia, Leptotene and Zygotene spermatocyte markers (DEGs identified in Fig. S1F) in indicated cell types from a 5 Y testis or the remaining samples (other).

Analysis of later stages of germ cell development further substantiated accurate reflection of in vivo processes, transition of pluripotency to spermatogenic programs, and puberty-induced meiotic progression. For example, many M.pSpg and T1.pSpg coexisted in 15-18 wpf testes (Fig. 1E, Table S4) where they tended to localize to central and peripheral region of the tubules, respectively (Fig. 1E)^17,26^. The proportion of M.pSpg gradually decreased and, by 24 wpf, were almost completely replaced by T1.pSpg by 24 wpf (Fig. 1E-G). Although T1.pSpg were present in 35 wpf testis (33.3% of all germ cells), they were dramatically decreased in number in 4 month old (4 M) postnatal testes (2.97%), largely replaced by (undifferentiated) spermatogonia (Spg) (Table S4). This population persisted until 12 year old, when spermatogenesis is largely completed as reflected by emergence of round spermatids (cell types 15-18, Std1-4). Notably, the T1.pSpg to Spg transition was characterized by further upregulation of spermatogonial marker genes (e.g., *UTF1*, *RET*, *TSPAN33*, *MAGEA4*, *EGR4*) and downregulation of a much smaller number of genes (e.g., *GAGE12H*, *HELLPAR*, *NANOS2*, *XIST*). Cells appear largely quiescent during this transition (i.e., lack of expression of *MKI67*, *UBE2C*, *TOP2A*), suggesting that the spermatogonial program was reinforced during the T1.pSpg to Spg transition (Fig. 1D, E, S2E). The spermatogonial state is followed by two sequential states of differentiation (early differentiating spermatogonia [Diff.Spg.E] and late differentiating spermatogonia [Diff.Spg.L]). Diff.Spg.E along with Spg, were the dominant population in 4 month-old testis. Diff.Spg.E were characterized by downregulation of a subset of spermatogonial markers (e.g., *UTF1*, *MORC1*, *TSPAN33*, *PIWIL4*, *EGR4*) without overt meiotic gene upregulation (Fig. S2F). Diff.Spg.L and subsequent meiotic spermatocytes appeared primarily in 12 Y testis, suggesting that onset of puberty triggers further differentiation of Spg and meiotic progression (Table S4). Interestingly, we noted a small number of germ cells entering into Diff.Spg.L and early meiotic phase (Leptotene) in 5 year-old testis (Fig. 1H, I), consistent with the previously reported abortive spermatogonial differentiation in prepubertal boys^27^. Germ cells in 12 year-old testis underwent different phases of meiosis (*PRDM9*^+^ pre.Leptotene [cell type 9], *SYCP1*^+^*DMRTC2*^+^ Leptotene [cell type 10], *SPACA1*^+^*CTAGE1*^+^ Zygotene [cell type 11], *OVOL1*^+^*SPATA8*^+^ Pachytene [cell type 12], *CCDC110*^+^*CMTM2*^+^ Diplotene [cell type 13], *TJP3*^+^*H1FOO*^+^ 2° Spermatocyte [cell type 14] and spermatids (Std) constituting 4 distinct cell types (*CCDC168*^+^*SPACA3*^+^ Std1 [cell type 15], *GSG1*^+^*TEX54*^+^ Std2 [cell type 16], *PRM1*^+^*TNP1*^+^ Std3 [cell type17], *TSSK6*^+^*SMCP*^+^ Std4 [cell type 18]) (Fig. 1D)^15,28–30^. By generating a comprehensive cell atlas of human male germline spanning the migratory PGC stage to the completion of spermatogenesis, our findings will serve as a key reference for understanding normal human male gametogenesis and neoplastic transformation, leading to seminoma and spermatocytic seminoma, which occurs in older men^9^.

### Single cell transcriptional landscape of seminoma and its tumor microenvironment

We next set out to determine the single cell transcriptomic landscape of seminoma tissues. As seminoma is characterized by its heavy immune cell infiltrate, we characterized both seminoma and additional cell types within the TME. For this analysis, we obtained seminoma tissues from two donors (Se1, Se2, Methods). Histologic sections revealed lobules of large proliferative polygonal cells with clear cytoplasm, large nuclei and prominent “cherry red” nucleoli separated by thin fibroconnective and vascular tissues that were heavily infiltrated by lymphocytes, features characteristic of pure seminoma (Fig. 2A, S3A)^9^. Immunofluorescence studies showed strong and uniform expression of *SALL4*, *NANOG*, *SOX17* and *TFAP2C* in both samples, further corroborating the diagnosis (Fig. 2B, S3B). A portion of these samples were dissociated and subjected to single cell transcriptomics. After filtering out low quality cells, a total of ∼22,615 cells (∼1,800 median genes/cell) were captured for downstream analyses (Fig. S3C). Overall, the two seminoma samples showed good concordance in cluster, cell cycle distribution and transcriptome (Fig. S3D-F). By projecting expression of known marker genes on the UMAP plot, we identified 7 cell types (cell type 1, *NANOG*^+^*TFAP2C*^+^ seminoma; cell type 2, *CD2*^+^GZMK^+^ T/NK cells, cell type 3, *LYZ*^+^*CD68*^+^ macrophages; cell type 4, *CD79A*^+^ B cells; cell type 5, *PLAC8*^+^ dendritic cells (DCs); cell type 6, *TAGLN*^+^ fibroblasts; cell type 7, *VWF*^+^ endothelial cells (Fig. 2C-E, S3G).

**Figure 2.**
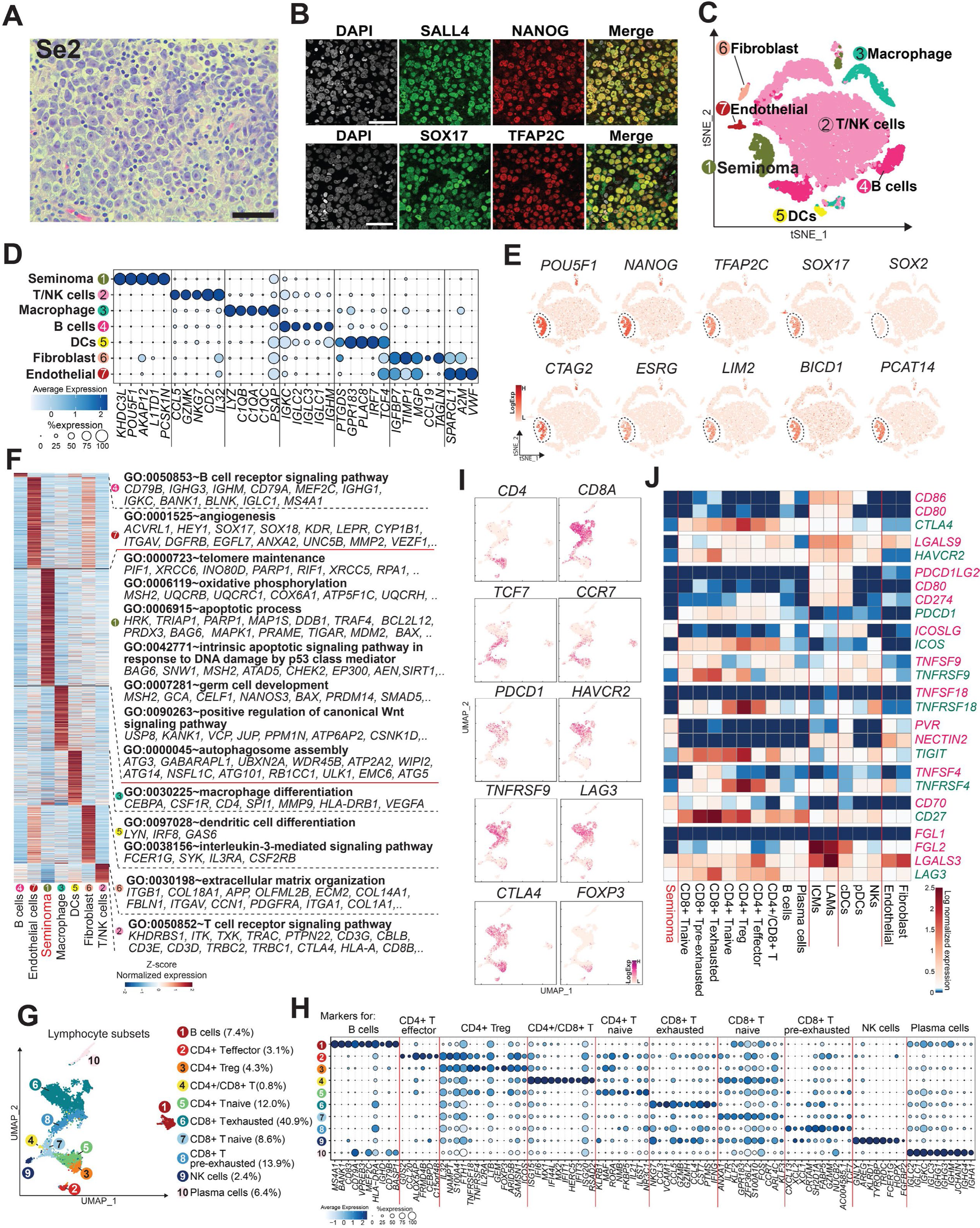
Human seminoma cell transcriptomic landscape at single-cell resolution. **(A)** Hematoxylin and eosin (H&E) staining of a section of seminoma sample (Se2). Scale bar, 100 μm. **(B)** IF of seminoma sections (Se2) for indicated proteins merged with DAPI staining. SALL4 and NANOG are pluripotency-associated proteins and SOX17 and TFAP2C are PGC specifier proteins. Scale bar, 50 μm. **(C)** tSNE plot of all seminoma cells based on computationally aggregated scRNA-seq data obtained from two seminoma samples (Se1, Se2). Different cell clusters were annotated based on marker genes and DEGs (Fig. 2D, E, S3G) and colored accordingly. **(D)** Dot plot of the top five DEGs by multi-group comparisons. Color code indicates expression, and circle size defines the portion of cells in certain cluster expressing the corresponding gene. Gene expression was log normalized and then Z-score transformed. **(E)** Expression of representative marker genes projected on the tSNE plot to define seminoma cells. **(F)** Heatmap showing the pseudo-bulk expression of DEGs among different cell types defined in (C). Representative genes and key GO enrichments are shown (right). **(G)** UMAP embedding showing the sub-clustering of lymphocyte lineages. T/NK cells and B cells in (C) are subsetted, then re-clustered. Cell types were annotated on the basis of marker genes and DEGs as (H, I). **(H)** Top ten DEGs obtained by multi-group comparisons in lymphocytes. Color code indicates pseudo-bulk expression, and circle size denotes the proportion of cells expressing the gene in indicated cell types. **(I)** Key markers of T cell exhaustion and immune checkpoint regulators projected on UMAP as defined in (G). **(J)** Pseudo-bulk expression of indicated cell types of seminoma TME for paired ligands (red) and receptors (green) associated with immune checkpoints. Expression is log2 (CPM+1) normalized. Color code indicates relative expression levels. ICMs, inflammatory cytokine-enriched macrophages; LAMs, lipid associated macrophages; cDCs, classical dendritic cells; pDCs, plasmacytoid dendritic cells; NKs, Natural killer cells.

Consistent with previous studies, comparative DEG analysis of seminoma cells revealed upregulation of transcription factors related to pluripotency (e.g., *NANOG, POU5F1, PRDM14, SALL4, TFCP2L1*) or germ cell specification (e.g., *TFAP2C*, *SOX17*), reminiscent of PGCs (Fig. 2D-F, S3G). Accordingly, these genes were enriched with GO terms such as “germ cell development”. Moreover, seminoma cells exhibited upregulation of a number of p53-target genes (e.g., *MDM2*, *AEN*, *SIRT1*, *BAX*) enriched with related GO terms (e.g., “apoptotic process” and “intrinsic apoptotic signaling pathway in response to DNA damage by p53 class mediator”) consistent with a wild-type p53-mediated pro-apoptotic state in seminoma (Fig. 2F, Table S5)^31,32^.

Within the TME, we found that fibroblasts express *TAGLN*, *ACTA2* and *FAP*, markers for cancer-associated fibroblast (CAF) (Fig. S3G)^33^. Sub-clustering analyses of tumor infiltrating lymphocytes (cell types 2 and 4) also revealed heterogeneity, with a dominance of CD8^+^ T cells with features of exhaustion (Fig. 2G-I, S3H, Table S6)^34,35^. Accordingly, immune checkpoint genes were widely expressed in T cells whereas many of their ligands were expressed on macrophages and classical (c)DCs (Fig. 2I, J). Sub-clustering of professional antigen presenting cells (cell types 3 and 5) revealed *C1QC*^+^*CHIT1*^+^*APOC1*^+^ lipid-associated macrophages (LAMs)^36–38^, *IL1B*^+^*CXCL8*^+^ inflammatory cytokine-enriched macrophages (ICMs)^36,38^, *BATF*^+^ cDC1s (cDC1s), *CD1C*^+^ cDC2, *IL7R*^+^*CD40*^+^*CCL19*^+^ activated/mature cDCs (A/M-cDCs and *TCF4*+ plasmacytoid DCs (pDCs) (Fig. S3I-L, Table S7)^39^. T cells with predominant features of exhaustion and expression of various immune checkpoint molecules support an overall immune-suppressive TME^40,41^.

### Comparison of transcriptomes of seminoma cells with normal human male germ cells

With single cell transcriptomes for normal human male germ cells and seminoma cells in hand, we next set out to map seminoma cells to the developmental coordinates of normal human male germ cells. When simultaneously projected onto a diffusion map, a short branch of seminoma cell trajectory bifurcated from the normal germ cell trajectory at the PGC.E/PGC.L/M.pSpg stage (Fig. 3A, B, S4A). Hierarchical clustering and Pierson’s correlation coefficient analyses showed the closest resemblance of seminoma cells to PGCs.E (Fig. 3C, S4B). We previously established an in vitro model of pre-migratory stage primordial germ cells (a.k.a., primordial germ cell-like cells [PGCLCs]) using human iPSCs^17,42^. Consistently, seminoma also exhibited the closest transcriptional similarity to PGCLCs (Fig. 3D, S4C). Seminoma cells and PGC(LC)s/M.pSpg shared a number of markers including pluripotency-associated (e.g., *POU5F1*, *NANOG*, *UTF1*, *L1TD1*, *KHDC3L*) and germ cell specifier genes (e.g., *PRDM1*, *PRDM14*, *TFAP2C*, *SOX15*, *SOX17*, *NANOS3*)^18,42–44^, some of which were over-expressed in seminoma cells compared to PGCs (e.g., *NANOG*, *KHDC3L*, *PRDM14*) (Fig. 3E-G, S4D-F). Of note, unlike PGCs.E/PGCs.L, seminoma cells do not express *DAZL*, *DND1*, or *DDX4*, markers normally upregulated in germ cells upon gonadal colonization^18,42,45^. However, they do express markers of PGCLCs that are not expressed in PGCs.E (e.g., *BICD1*, *ANGPTL4, PRDX2*) suggesting that they might have some pre-migratory features akin to PGCLCs (Fig. 3E-G, I, S4H).

**Figure 3.**
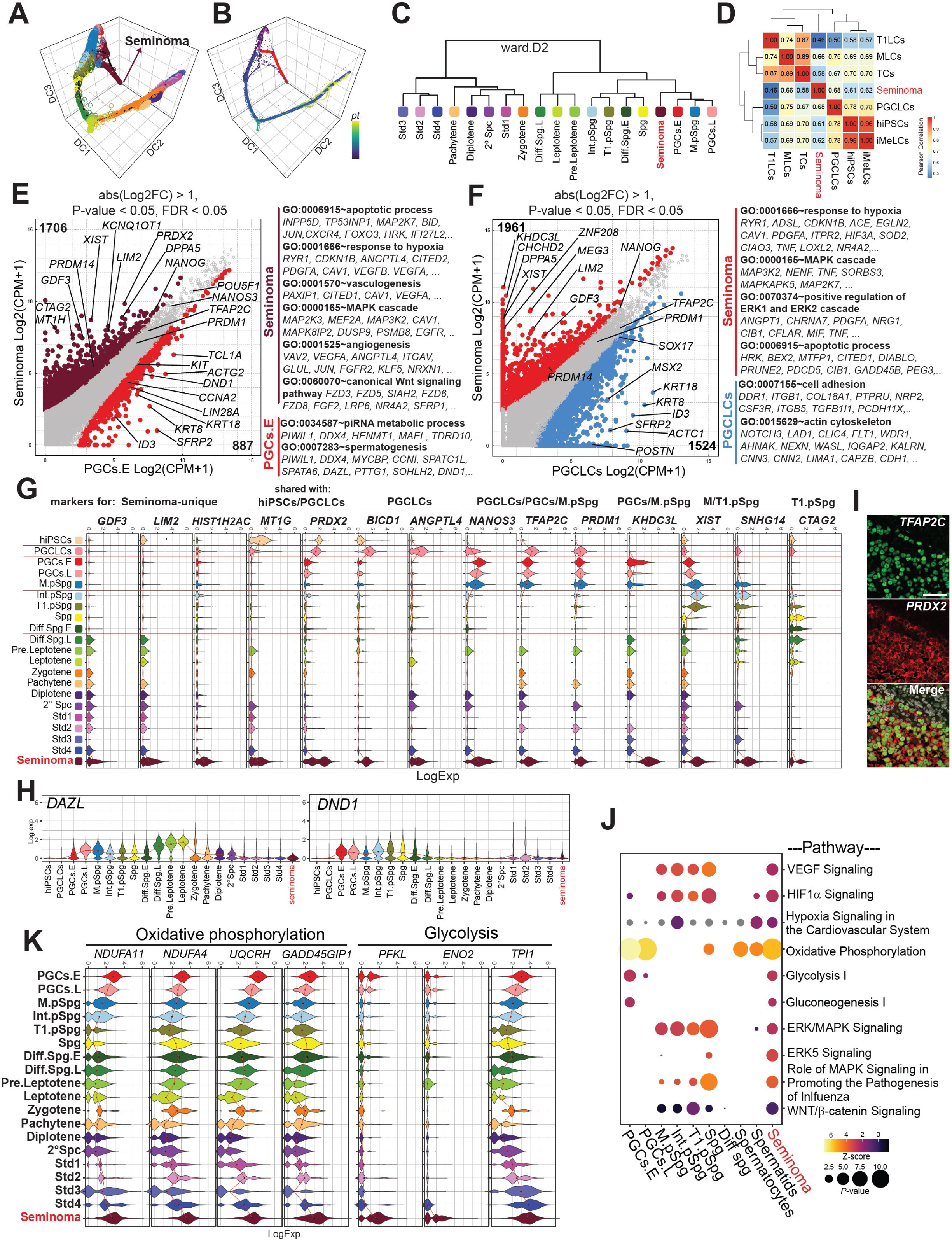
Normal and abnormal developmental trajectories of human male germline using single cell transcriptomes. **(A, B)** 3D DiffusionMap projections of annotated cell types (**A**) and pseudotime **(B).** Cell types are defined in Fig.1A and 2C. The bifurcated pseudotime trajectories show seminoma cells (highlighted in red) diverged from normal germline development. Cells are colored according to cell types in (**C**). **(C)** Unsupervised hierarchical clustering of normal male germ cell types and seminoma using Euclidean distance of pseudo-bulk transcriptomes of indicated cell types. Top 2000 variable genes were used. **(D)** Heatmap showing Pearson correlation of seminoma and cell types generated during male germ cell induction from iPSCs^17^. T1LCs, T1.pSpg-like cells; MLCs, M.pSpg-like cells; TCs, transitional cells; PGCLCs, PGC-like cells; iMeLCs, incipient mesoderm-like cells. **(E, F)** Scatter plots comparing the averaged gene expression values between seminoma and PGCs.E (E) or seminoma and PGCLCs (F). Expression is normalized by log2(CPM+1). DEGs (absolute log2 fold change > 1, p-value < 0.05 and FDR < 0.05) are highlighted in colors. Representative genes and their GO enrichments for DEGs are shown on the right. **(G)** Violin plot showing expression of representative marker genes of the indicated normal male germ cells that are also expressed in seminoma. Gene expression is log normalized. **(H)** Violin plot showing expression of *DAZL* and *DND1* in the indicated cell types. **(I)** Co-in situ hybridization (ISH)/IF images of a seminoma tissue section for TFAP2C (green) and *PRDX2* (red) merged with DAPI staining (white). Scale bar, 50 μm. **(J)** Ingenuity signaling pathway enrichment analysis of normal germ cells and seminoma. **(K)** Violin plot showing expression of genes related to oxidative phosphorylation and glycolysis in seminoma and normal male germ cells.

Despite the overall similarity to PGCS.E/PGCLCs, seminoma cells also express markers that are not expressed in these cells. For example, seminoma cells share some markers with iPSCs (e.g., *MT1G*), and T1.pSPg/Spg (e.g., *CTAG2*, *MAGEB2*) (Fig. 3G, S4F, G). We also identified genes that are highly expressed in seminoma, but either not expressed or only weakly expressed in iPSCs or any germ cells that were analyzed (e.g., *GDF3*, *LIM2*, *HIST1H2AC, ESRG*) (Fig. 3G, S4A, H), suggesting that they might be useful for diagnosis and/or disease monitoring. Moreover, pairwise transcriptomic comparison of seminoma cells to PGCs (PGCs.E or PGCs.L), or to M.pSpg revealed upregulation of a number of genes in seminoma cells associated with MAPK/ERK signaling, hypoxia/angiogenesis, canonical Wnt signaling and the apoptotic process, any of which might be involved in seminoma progression and their transformation from normal PGCs (Fig. 3E, F, J, S4D, E, I). Like PGCs, seminoma cells activate genes associated with both oxidative phosphorylation and glycolysis, suggesting a unique metabolic state (Fig. 3J, K)^22^. However, seminoma cells tend to express even higher levels of glycolytic genes (enriched with GO terms, “glycolytic process”) (Fig. S4I), which might reflect their adaptation to hypoxic condition within the TME.

Sub-clustering of seminoma cells revealed intra-tumor heterogeneity and inter-patient variability within 5 subclusters (Fig. S5A-D), which aligned along the pseudotime trajectory from proximal to distal orientation. Progressive upregulation was noted along pseudotime for some genes related to pluripotency/germ cell specification, chromosome 12p (e.g., *CCND2*, *KDM5A*, *KRAS*) or angiogenesis (e.g., *EMC10*, *ANGPTL4*) (Fig. S5E). Accordingly, clusters distant from PGCs showed upregulation of genes whose expression deviated from that of normal germ cells and were enriched with Ingenuity pathways or GO terms related to angiogenesis, hypoxia, or ERK/MAPK signaling, which might reflect seminoma cell evolution (Fig. S5F, G). On the other hand, cluster 1, most proximal to PGCs in vivo, showed expression of genes related to oxidative phosphorylation akin to PGCs.E.

### Chromatin accessibility in human fetal male germ cells

To better define the cellular identity of seminoma, we utilized a single-nucleus muti-omic approach that allow for concomitant genome-wide nuclear mRNA expression analysis (scRNA-seq) and chromatin accessibility (ATAC) analysis (ATAC-seq) in both normal and transformed tissue. Although access to human fetal tissues, particularly those at the 1^st^ trimester is limited, we were able to obtain a fetal testis from a 2^nd^ trimester fetus (21 wpf) for multi-omics analysis (Fig. S6A-C). Analysis of gene expression and chromatin accessibility individually or in combination allowed us to confirm distinct cell types (Fig. S6D) previously identified in solo scRNA-seq (Fig. S1C)^17^. A number of differentially accessible regions (DAR) were identified across these cell types, suggesting that regulation of the chromatin landscape was cell-type specific (Fig. S6E, F). For example, germ cells showed strong chromatin accessibility peaks at the proximal and/or distal regions of *TCL1A* and *NANOG*, likely representing promoter and enhancer regions, respectively (Fig. S6G). These cell types exhibited unique DEGs based on nuclear mRNA expression (Fig. S6H), which were consistent with previous solo scRNA-seq data (Fig. S1D)^17^. We also calculated gene activity scores based on accessibility at promoter regions to infer gene activity and compare them across cell types, allowing us to generate a list of genes with differential gene activity scores (DGAS). This analysis also identified genes unique to each cell type, which were largely in agreement with mRNA expression (Fig. S6I). For example, both DEG and DGAS analyses revealed *DCAF4L1* in germ cells, *KDR* in endothelial cells, and *INHA* in Sertoli cells (Fig. S6H, I).

To better define cell states, we cross-referenced TF DNA-binding motif enrichment within DAR and DEGs and DGAS to identify TFs that were both highly expressed and have access to their target binding motifs, thereby promoting target gene transcription (herein designated as active TF analysis). This analysis identified known TFs with germline expression/function (e.g., *KLF5*, *TFAP2C)* along with previously uncharacterized TFs (e.g., *REST*, *ZNF682*, *TCF3*) (Fig. S6K)^43,44,46^. Enriched TFs in other cell types also revealed well-recognized genes, further highlighting the validity of this approach (e.g., *TCF21* and *LHX9* in gonadal stroma, *GATA4* and *SOX9* in Sertoli cells)^20,47,48^. Sub-clustering of germ cell populations in UMAP revealed two distinct cell types (Fig. S7A). UMAP projection of nuclear mRNA expression and gene activity scores for unique markers revealed that these cells represent M (marked by *POU5F1*, *NANOG*, *TFAP2C*, *PRDM1*) and T1 pSpg (marked by *PIWIL4*, *MORC1*, *CTAG2*), which progressed from M to T1.pSpg by pseudotime analysis (Fig. S7B-D). Although Int.pSpg were not identified by this sub-clustering analysis, likely due to its sample size, *ASB9*, a marker of Int.pSpg highlighted the cells at the junction of M.pSpg and T1.pSpg (Fig. S7C)^17^. Motif enrichment analysis of DAR also revealed that motifs for the well-known pluripotency-associated TFs (e.g., *POU5F1*, *NANOG*) and germ cell specifier TFs (e.g., *TFAP2C*, *SOX17*) were among the most enriched in M.pSpg but that such enrichment was diminished as they progress towards T1.pSpg (Fig. S7D, E). Open chromatin peaks unique in M.pSpg or T1.pSpg were primarily located within non-promoter regions, consistent with the notion that these peaks represent enhancers that are often cell type specific and responsible for cell fate transition (Fig. S7F, G). Accordingly, inspection of *TFAP2C* or *NANOG* loci revealed unique distal peaks in M.pSpg that might serve as cell type specific enhancers (Fig. S7H). Although TF motif enrichment in M.pSpg was generally in agreement with expression, it exhibited some discordancy in T1.pSpg; some TFs with enriched motifs were either not expressed (e.g., *HOXA9*, *ELF4*) nor more highly expressed than in M.pSpg (e.g., *DMRT1*). On the other hand, enrichment was not observed in many spermatogonia specific TFs (e.g., *EGR4*, *ID1*, *ID2*, *ESX1*).

### Chromatin accessibility in the seminoma microenvironment

Using the multiome approach, we next set out to define the gene expression and chromatin accessibility landscape of the same seminoma tissue sample used for solo scRNA-seq (Se2). The combination of scRNA-seq and ATAC-seq data projected on UMAP revealed 7 distinct cell types identical to those previously defined by solo scRNA-seq (Fig. 2C, S8A). The majority of seminoma cells appeared in cluster 5, which differed the most significantly from normal germ cells (Fig. S5B, C). There was a high degree of concordance of gene expression and gene activity scores (Fig. S8B, C). Annotated cell types revealed cell-type specific DEGs and DGAS largely in agreement with DEGs obtained in solo scRNA-seq (Fig. S8D, E), where seminoma cells exhibit high gene expression and gene activity scores for pluripotency-associated and germ cell specifier genes (e.g., *NANOG*, *TFAP2C*) (Fig. S8C-E). Active TF analysis also revealed *PLAG1*, *SP4*, *REST*, *TFAP2C*, *ZNF682* and *KLF11* as seminoma specific active TFs (Fig. S8G), among which *REST*, *TFAP2C* and *ZNF682* were shared with germ cell specific active TFs (Fig. S6K), highlighting the similarities between seminoma and human fetal germ cells.

### Integrative multi-omics analysis of seminoma cells and human fetal germ cells

To compare the transcriptional and chromatin signatures between seminoma cells and human fetal germ cells, we integrated their multi-omics data and projected them onto UMAP. This integrative analysis yielded heterogeneity of both seminoma and normal germ cells with two seminoma cell clusters (C1, C2) and two germ cell clusters (C3, C4) (Fig. 4A-C). Two germ cell clusters (C3, C4) were readily identified as M.pSpg and T1.pSpg, respectively based on marker gene expression/gene activity scores (e.g., M.pSpg marker, *POU5F1* in C3 and T1.pSPg marker, *MAGEC2* in C4) (Fig. 4D), the former of which shows a higher correlation with seminoma cells (Fig. 4C). Despite overall similarities, there were notable differences between seminoma cells and M.pSpg (e.g., both gene expression and gene activity score for *NANOG* and other genes on chromosome 12p, many of which are associated with pluripotency and were substantially higher in seminoma cells than in M.pSpg) (Fig. 4D, E). Accordingly, the prediction of cis-regulated genes from DAR using GREAT analysis revealed stem cell-related GO terms enriched in seminoma cells compared to M.pSpg (e.g., “regulation of stem cell population maintenance) (Fig. 4F). DEG and DGAS analyses comparing seminoma cells (C1, C2) and germ cells (M.pSpg, T1.pSpg) revealed that seminoma cells expressed GOs associated with stem cells and germ cell development (e.g., “somatic stem cell population” or “germ cell development”) along with GOs associated with neoplastic features (“angiogenesis”, “MAPK cascade”), consistent with findings obtained by solo scRNA-seq (Fig. 4G, H). Finally, active TF analysis further confirmed that seminoma cells showed enrichment of many key germ cell specifier genes (*KLF4* and *TFAP2C* in C1, *PRDM1*, *NANOG* in C2 (Fig. 4I). Consistently, similar to M.pSpg, seminoma had proximal peaks flanking the transcription start site (TSS) of *NANOG* (−3.0kb, −2.2kb, +1.6kb), albeit at a higher intensity (Fig. 4J). Notably, seminoma cells also had multiple unique peaks at distal regions that were not seen in M.pSpg, suggesting that they might serve as seminoma specific enhancers that contribute to the over-expression of these core germ cell genes (Fig. 4J). Active TF analyses also identified previously uncharacterized TFs enriched in seminoma cells including *ENO1* and *KLF11* in C1 and *CUX1*, *CUX2*, *BACH1*, *ZNF75D* in C2, which might exert a unique function in seminoma and warrant future investigation (Fig. 4I).

**Figure 4.**
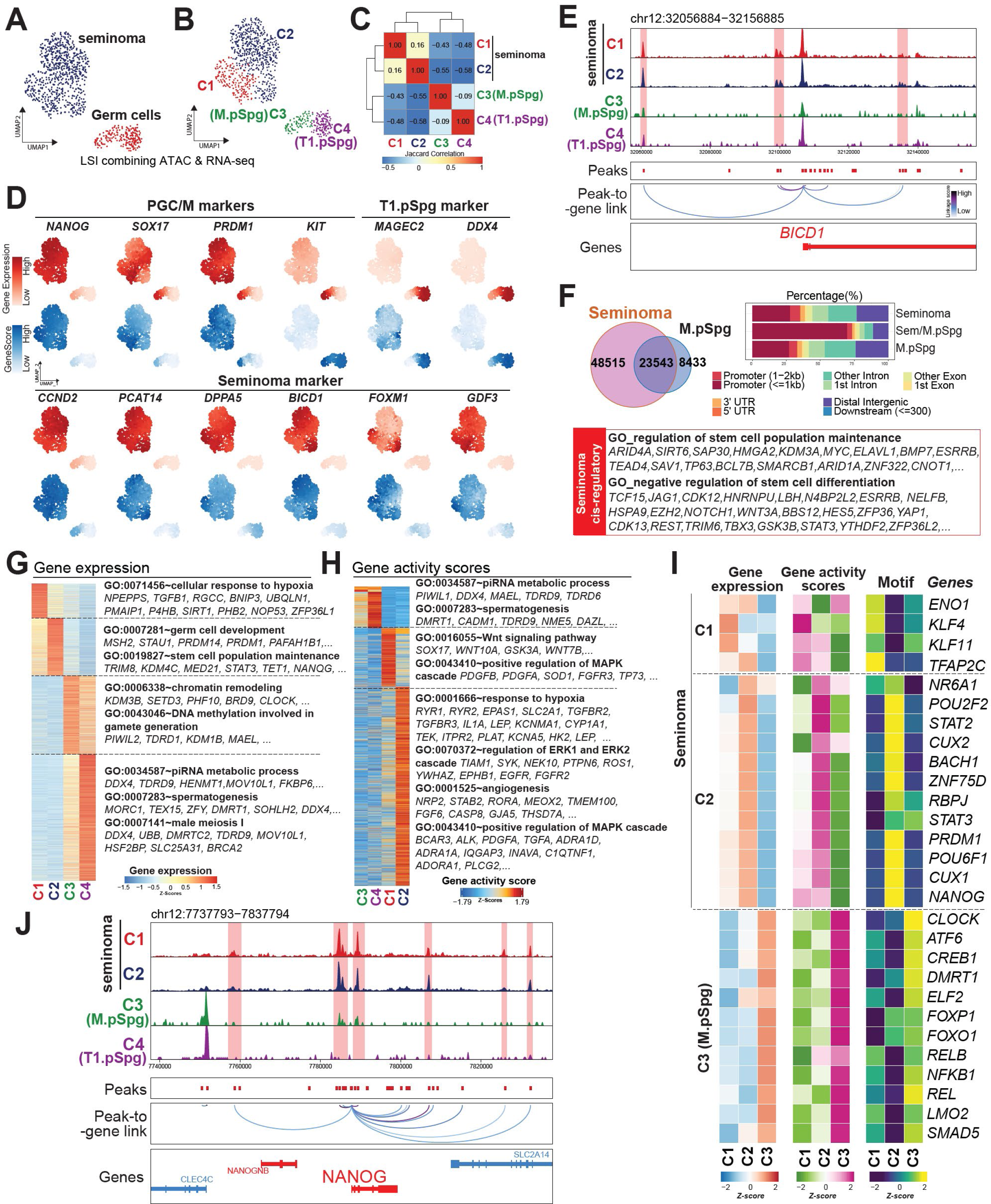
Comparison of seminoma and fetal germ cells by single cell multiomics. (**A, B**) Latent semantic indexing (LSI)-UMAP projection of combined single cell ATAC and RNA-seq data. Cells are colored according to cell types (A) or cell clusters (B). Cell types are defined in Fig. S6D and S8A. Cell clusters, C3 and C4 represent M.pSpg and T1.pSpg, respectively based on expression of marker genes as in (D). **(C)** Jaccard correlation of chromatin accessibility among different cell clusters. **(D)** Projection of markers of PGCs/M.pSpg, T1.pSpg and seminoma defined by scRNA-seq (Fig.2E, 3E-G) for gene (RNA) expression and gene activity scores onto LSI-UMAP embedding. **(E)** Genome browser screenshot of *BICD1* locus. Each track shows pseudo-bulk ATAC-seq signal for indicated cell types. Note the dynamic distribution of peaks at the distal regulatory regions (highlighted in red). Peak-to-gene linkage shows strength of correlation between chromatin accessibility and gene expression. **(F)** Genomic annotation of open chromatin regions identified in seminoma cells (C1 and C2) and M.pSpg (C3) are categorized in peaks unique to seminoma cells or M.pSpg or overlapped between them (top left). Stacked barplots show genomic distribution of open chromatin peaks in seminoma cells and M.pSpg (top right). GREAT GO enrichment analysis inferring cis-regulation by distal intergenic open chromatin regions unique in seminoma cells. Representative putative target genes and key GO enrichments are shown. (**G, H**) Heatmap showing the averaged values of DEGs (G) or differential gene activity scores (DGAS) (H) identified from a multi-group comparison among cell clusters. Gene expression and gene activity scores are Z-score normalized. In each heatmap, values are Z-score normalized. Representative genes in each GO category are shown. **(I)** Integrated analysis of gene expression (RNA), gene activity scores and motif enrichment (Motif). Genes whose values in all of the above categories are significantly higher in one cell cluster over the other clusters are shown as “active” transcription factors. **(J)** Genomic browser screenshot of *NANOG* locus as in (E).

### Seminoma exhibits transposable element (TE) expression reminiscent of PGCs

Early embryonic development, including that in the early germline, is often accompanied by activation of various TEs that are expressed in a stage/cell type specific manner^17,49–51^, partly as a result of ongoing DNA demethylation^52^. Notably, similar classes of TEs are upregulated in PGCs/M.pSpg and seminoma^53–55^. However, as the expression dynamics of TEs in seminoma is only partially understood, we next used our scRNA-seq data to characterize TE expression in seminoma and cells within the TME. Dimension reduction analysis of genome-wide TE expression projected on tSNE revealed separation of cell types, supporting cell-type/stage specific TE expression (Fig. 5A). Moreover, consistent with its overall hypomethylated genomic status^53^, overall TE expression levels were higher in seminoma than in TME cells (Fig. 5B), with L1 (LINE), Alu (SINE) and ERV families (LTR) being particularly enriched in seminoma cells (Fig. 5C). Moreover, the upregulated TE in seminoma cells were highly enriched in evolutionary young LTR TE (e.g., LTR7/HERVH-int, LTR17/HERV17-int [primate-specific ERV1]; LTR5-Hs, HERVK-int [hominoid-specific ERVK]) (Fig. 5D-F, Table S8).

**Figure 5.**
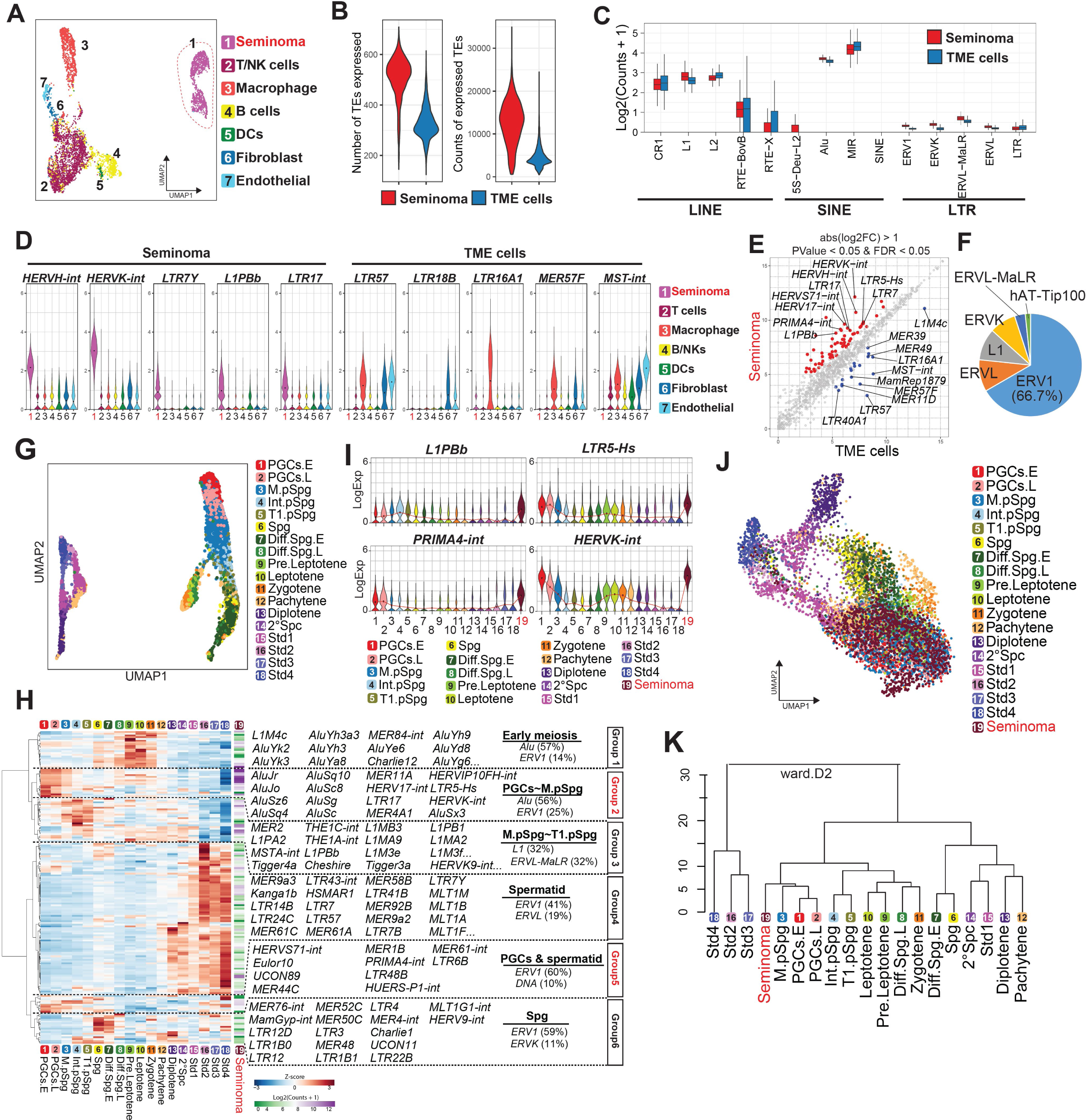
TE expression dynamics in seminoma and human male germline. **(A)** UMAP plot of all cells derived from seminoma tissues by using TE expression of aggregated scRNA-seq data. Cells are colored according to cell types identified in Fig. 2C. **(B)** Violin plot showing the number of TE features (left) and total TE counts (right) in seminoma cells (red) and the remaining cells of their tumor microenvironment (TME cells) (blue). **(C)** Box plot showing the log2 normalized averaged expression of TEs of different families in seminoma (red) or TME cells (blue). Expression is normalized by overall size of TEs per library. **(D)** Violin plots showing averaged TE expression that are unique to seminoma (left) or other TME cell types (right). **(E)** Scatterplot of pair-wise comparison between seminoma and TME cells. Differentially expressed TEs are shown in colored dots (defined as absolute log2Fold change > 1, P value < 0.05, FDR < 0.05). Expression value is normalized as log2(CPM+1). Key TEs are highlighted. **(F)** Pie chart showing the proportion of TE subfamilies among differentially expressed TEs upregulated in seminoma as in (E). **(G)** UMAP plot of male germ cells clustered by using TE expression of aggregated scRNA-seq data. Cells are colored based on cell types identified in Fig. 1A. **(H)** Heatmap showing the averaged expression values of the top 100 variably expressed TEs across all human male germ cells. Expression is Z-score normalized. Expression of respective TEs in seminoma cells are shown on the right, which are normalized by log2 (counts +1). These TEs are clustered largely into 6 groups according to their expression dynamics. Group, TEs upregulated in early meiotic stages; Group 2, TEs upregulated in PGCs to M.pSpg stages; Group 3, TEs upregulated M to T1.pSpg stages; Group 4, TEs upregulated in spermatid stages; Group 5, TEs upregulated in PGCs and spermatid stages; Group 6, TEs upregulated in Spg. Top 2 TE subfamilies represented in each group are shown with their percentages. **(I)** Violin plot showing examples of TEs co-expressed in PGCs/M.pSpg and seminoma cells. **(J)** UMAP embedding showing both seminoma cells and human male germ cells. Note that seminoma cells overlap with PGCs.E/PGCs/L and M.pSpg. **(K)** Hierarchical clustering (Ward.2 method using Euclidian distances) of the respective cell types (averaged expression values) using the 200 most variably expressed TEs.

To establish references for seminoma cells, we next analyzed TE expression dynamics during male germline development (Fig. 5G, H). Consistent with our previous study using a more limited data set^17,49^, this analysis revealed dynamic stage-specific expression of TEs. For example, TEs upregulated in prospermatogonia (Group 3) were enriched with L1 (LINE) (Fig. 5H). Spg and diff.Spg.E express unique sets of LTR TEs (e.g., LTR12 family) consistent with previous studies^30^. PGCs and M.pSpg (Group 2 and 5) were characterized by high expression of Alu (SINE) and primate- or hominoid-specific LTR TEs (e.g., *LTR5-Hs, HERVK-int, LTR17, HERV17-int, PRIMA4-int*), reminiscent of seminoma cells (Fig. 5H, I). Consistently, these cells were clustered with seminoma cells in UMAP and by hierarchical clustering analysis (Fig. 5J, K). Given the emerging role of TEs as cis-regulatory elements of early embryonic and germline development, these findings suggest that the gene regulatory network orchestrated by TEs in seminoma might be operated in a manner similar to PGCs.

### Isochromosome 12p triggers KRAS expression and MAK kinase activity in PGCLCs

The vast majority of seminoma carry the pathognomonic marker chromosome, isochromosome 12p [i(12p)], which bears two additional copies of the short arm of chromosome 12 (12p) (Fig. S9A)^6,8,9^. Indeed, copy number variation (CNV) analysis using seminoma and normal germ cell transcriptomes revealed overrepresentation of the 12p region (Fig. S9B, C). 12p contains key genes involved in both germ cell development (e.g., *DPPA3* and *NANOG*) and carcinogenesis (e.g., *KRAS*, *CCND2*). However, the causal relationship of i(12p) in driving seminoma remains unclear, in part due to the lack of an appropriate model system.

Individuals with Pallister Killian syndrome (PKS), a multi-organ system developmental diagnosis, are mosaic for i(12p), representing an opportunity to explore the impact of i(12p)^19^. Given our scRNA-seq data showing the similarities of PGCLCs to seminoma cells (Fig. 3D), and our ability to generate human PGCLCs from iPSCs, we created PGCLCs with or without i(12p) by reprogramming isogenic iPSCs from PKS fibroblasts, providing us with the unprecedented ability to test the dosage effect of i(12p) on the PGC transcriptome (Fig. 6A). We established iPSCs from fibroblasts of three male patients with PKS [PK12, PK19, PK44, 65%, 100%, 96% bearing i(12p), respectively] (Fig. S9C, D). From two patients (PK12, PK19), we successfully established iPSC lines with and without i(12p) allowing us to evaluate its effect in an isogenic setting. iPSCs with or without i(12p) grew at a similar rate and had similar levels of pluripotency marker expression (Fig. S9E, F). Moreover, hiPSCs with i(12p) formed teratomas, suggesting that i(12p) does not affect pluripotency (Fig. S9G). PGCLCs were readily induced from iPSCs bearing i(12p) without a significant alteration in either induction efficiency or expression of key germ cell markers as assessed by qPCR (Fig. 6B-D). Notably, however, when projected on the PCA axis or analyzed by hierarchical clustering, transcriptomes of PGCLCs showed clear segregation between those with and without i(12p) (Fig. 6E, F). Pairwise transcriptomic comparison of iPSCs or PGCLCs with and without i(12p) revealed DEGs, the majority of which were upregulated in those with i(12p) and were highly enriched in genes on chromosome 12p, consistent with a dosage effect caused by extra copies of chromosome 12p (Fig. 6G, H). The number of upregulated DEGs was higher in PGCLCs than in iPSCs, albeit with lower enrichment of genes on chromosome 12p, likely due to the secondary changes triggered by upregulated 12p genes (Fig. 6G, H). Notably, upregulated DEGs were enriched in GO terms such as “canonical Wnt signaling pathway” (e.g., *TCF7L2*, *FZD7*, *LRP6*), “angiogenesis” (e.g., *NRP2*, *ANGPT1*), “positive regulation of MAP kinase activity” (*PDGFA*, *KRAS*, *FGFR1*) (Fig. 6G, H), reminiscent of transcriptomic features of seminoma. In keeping with this pattern, PGCLCs with i(12p) showed higher Pearson correlation with the seminoma transcriptome than did those without i(12p) (Fig. 6I). The majority of genes (61.5%) commonly upregulated between seminoma (vs somatic infiltrates or vs PGCs.E) and PGCLCs with i(12p) [vs PGCLCs without i(12p)] were derived from chromosome 12p (e.g., *PDGFA*, *KRAS*, *FGFR1, CCND2*) and enriched with GO terms such as “positive regulation of MAP kinase activity” or “MAP cascade” (Fig. 6J-L). Thus, consistent with scRNA-seq and multiome analyses, these findings suggest that these genes will contribute to the overactivation of MAP kinase signaling in seminoma cells. Collectively, these data support the causative role of extra copies of 12p in shaping genetic programs that promote transformation of seminoma cells from normal germ cells.

**Figure 6.**
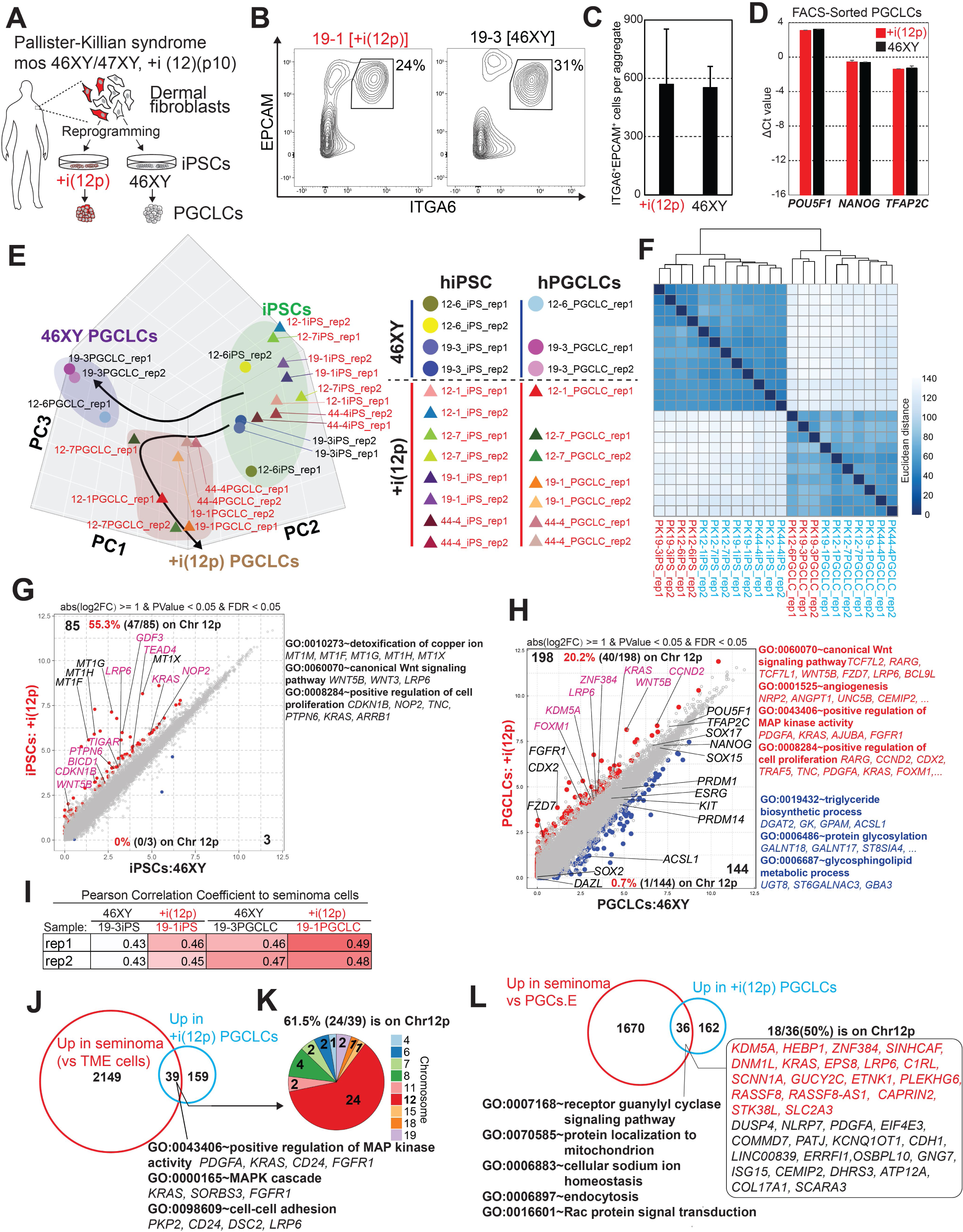
*In vitro* seminoma model using PGCLCs bearing isochromosome 12p. **(A)** Experimental design used to derive PGCLCs with or without i(12p) from isogenic iPSCs established from dermal fibroblast of patients with Pallister-Killian syndrome bearing i(12p) mosaicism. **(B)** FACS plots of day 5 PGCLC-containing aggregates induced from wild type iPSCs (46XY) or iPSCs with i(12p) [+(12p)] iPSCs. Percentages of PGCLCs labeled by EPCAM and ITGA6 are shown. **(C)** FACS quantification of the total number of PGCLCs (ITGA6^+^EPCAM^+^ cells) per aggregate induced from isogenic iPSCs with or without i(12p). Averaged values +SD from 3 biological replicates are shown. **(D)** qPCR quantification of key human PGC marker genes in cDNA libraries generated from FACS-sorted day 5 PGCLCs (ITGA6^+^EPCAM^+^) with and without i(12p). For each gene examined, the ΔCt from the average Ct values of the two housekeeping genes *ARBP* and *PPIA* (set as 0) are calculated and plotted. **(E)** Principal component analysis (PCA) of bulk transcriptomes of iPSCs and PGCLCs of indicated sample origins. Samples bearing i(12p) are highlighted in red. **(F)** Heatmap showing Euclidean distance and Hierarchical clustering (Ward.2 method) of indicated bulk transcriptome samples. Samples bearing i(12p) are highlighted in red. **(G, H)** Scatterplot comparing the averaged gene expression values of iPSCs (G) or PGCLCs (H) between indicated karyotypes. DEGs (highlighted as colored dots) are defined with absolute log2 fold change > 1, p-value < 0.05, FDR < 0.05. Expression value is normalized as log2(CPM+1). Enriched GO terms and representative genes are shown on the right. Percentages of genes on chromosome 12p (colored in purple) among DEGs are shown. **(I)** Pearson correlation of averaged expression values of indicated transcriptomes comparing to seminoma transcriptome. **(J)** Venn diagram showing overlapping genes (39 genes) between up-regulated DEGs in seminoma (versus TME cells as in Fig. 2F) and up-regulated DEGs in PGCLCs with i(12p) [versus isogenic control without i(12p) as in (H)]. Enriched GO terms and representative genes are shown at the bottom. **(K)** The proportion of chromosomal location of the overlapping genes. **(L)** Venn diagram showing overlapping genes between up-regulated genes in seminoma (versus PGCs, identified in scRNA-seq) and up-regulated DEGs in PGCLCs with i(12p) [versus isogenic control without i(12p) as in (H)]. Enriched GO terms and genes are shown with those on chromosome 12p highlighted in red.

Taken together, our comprehensive analysis of seminoma in comparison to normal germline highlights the genetic similarities and differences that might contribute to seminoma pathology and that may serve as future therapeutic targets.

## DISCUSSION

Human male germline development is a complex and temporally coordinated process. Deviation can result in infertility and germ cell tumor formation. To obtain insight into the development of seminoma and inform on malformations that can impact normal development and fertility, we employed scRNA-seq to characterize a wide-ranging and complete set of fetal, neonatal, and prepubertal testicular samples to generate a comprehensive and continuous lineage trajectory map of human male germ cells spanning from migrating PGC to the haploid spermatid stages.

Analyses revealed progressive yet somewhat asynchronized cell fate transitions that confer cellular heterogeneity in the testes. For example, the migrating PGC state (PGCs.E), characterized by high overall pluripotency and germline specifier gene expression, does not immediately proceed to the M.pSpg state upon gonadal colonization (occurring at 5-7 wpf), as some PGCs.E cells remain at 9 wpf. Likewise, transition of M.pSpg, characterized by modest downregulation of pluripotency-associated genes and upregulation of proliferation associated genes (etc. *MKI67*, *FOXM1*), to T1.pSpg occurs via an *ASB9*^+^ intermediate state (Int.pSpg) in an asynchronized fashion. The function of ASB9 in M.pSpg to T1.pSpg transition remains unclear but it might be involved in inhibition of cell growth through modulation of MAPK signaling^56,57^. Transcriptionally, the M-to-T1.pSPg transition is characterized by loss of proliferation, loss of pluripotency-associated and germline specifier genes and upregulation of spermatogonia-specific (spermatogenic) TFs. Interestingly, the expression of spermatogonia-specific TFs in T1.pSpg is generally lower than that of Spg and their chromatin accessibility is limited (Fig. S7E), suggesting that the spermatogenic program is not yet fully operative. Moreover, the concomitant loss of chromatin accessibility to pluripotency and germline specifier TFs characteristic of M.pSpg suggests that shutting off pluripotency, rather than spermatogenic transcriptional reprogramming is the hallmark of T1.pSpg. Our analysis also provided convincing evidence of abortive meiotic entry (i.e., leptotene stage) in a prepubertal boy, which was previously suggested only by histomorphologic observations^27^. The commonality of this phenomenon warrants further confirmation but supports the spermatogenic potential for prepubertal spermatogonia. The more robust meiotic entry (as seen in 12 Y testis) followed by completion likely dependents on puberty and an increase in local and systemic testosterone levels^15,58^.

With the comprehensive germ cell developmental map, we were able to pinpoint the position of seminoma cells in the normal human germ cell transcriptomic trajectory. Remarkably, this analysis revealed that seminoma cells resemble PGCs.E (migratory PGCs). The comparison of seminoma to iPSC-derived PGCLCs, representing pre-migratory PGCs, also revealed a high degree of similarity, supporting their potential use as an in vitro platform to dissect both normal and pathologic germ cell development. Specifically, seminoma exhibits low expression levels of genes that are typically upregulated after gonadal colonization (i.e., *DDX4*, *DAZL*) (Fig. S2)^13,45^. Notably, *DAZL* promotes germ cell commitment by suppressing the pluripotency program, and their absence can facilitate germ cell tumor formation^59,60^. Moreover, migratory/pre-migratory PGCs (i.e., PGCs.E, PGCLCs) show higher expression of pluripotency-associated genes than do gonadal germ cells (i.e., PGCs.L, M.pSpg). Likewise, seminoma cells exhibit enhanced pluripotency-associated TF activities by multiomics analysis (e.g., *KLF4*, *POU2F2*, *POU6F1*, *NANOG*) (Fig. 4I). While the molecular features associated with migratory/pre-migratory PGCs might confer a permissive state for seminoma formation and/or progression, elucidation of such molecular mechanisms await development of model systems as discussed below.

Despite similarities, seminoma cells clearly differ from normal PGCs. Under physiological conditions, human PGCs are unipotent and their conversion into pluripotency is exceptionally rare. However, seminoma cells often manifest overt pluripotency, as revealed by concomitant embryonal carcinoma and/or teratoma formation with the same clonal origin^61,62^. Our direct comparison of seminoma with PGCs revealed upregulation of various pluripotency-associated genes in seminoma cells that are either expressed higher in seminoma than in PGCs (e.g., *NANOG*, *PRDM14*) or are not highly expressed in any germ cells observed (e.g., *GDF3*, *DPPA5*). The molecular basis for the over-activation of pluripotency programs is unclear, but it might be controlled, in part, by unique epigenetic rewiring in seminoma. In support of this notion, we identified an open chromatin accessibility of the distal region of the *NANOG* loci in seminoma cells, but not in normal germ cells (Fig. 4J).

Our direct transcriptome comparison also reveals upregulation of a number of genes involved in apoptotic pathways, many of which are known targets of p53 (e.g., *TRIAP1, SIRT1, MDM2, BAX*)^32^. This finding is consistent with previous studies revealing either intact or hyperactivated p53 pathways in the majority of germ cell tumors^63–66^, a property that confers cisplatin sensitivity^31^. Another remarkable finding in our transcriptomic comparison is upregulation of genes related to MAPK/ERK activation in seminoma cells (Fig. 3E, F, J, 4H, S4D, E, S5F, G). Importantly, there are many genes related to MAPK/ERK signaling on chromosome 12p, which is often present in multiple copies in seminoma cells. We found that PGCLCs bearing i(12p) exhibited upregulation of key genes related to MAPK pathway (e.g., *KRAS*, *PDGFA*, *FGFR1*), highlighting the utility of the hPGCLC-based system for predicting seminoma genotype-phenotype correlation. Although PGCLCs with i(12p) exhibited more similarities to seminoma than wild-type PGCLCs (Fig. 6I), many genes that were upregulated in seminoma cells compared to PGCs were still not upregulated and PGCLCs with i(12p) did not exhibit overt growth increase, indicating that i(12p) is not sufficient for seminoma transformation (Fig. 6C, J, L). Interestingly, a PKS patient with a presumably germ cell origin pineal tumor was previously reported, although the commonality of these events remains unknown^67^. To determine signaling events driving tumor formation, future studies will include expression of activated *KIT* or *KRAS* mutations using the in vitro platform, as these mutations are frequently seen in seminoma, and can also activate downstream MAPK signaling^10,64^.

Through coordinated, unbiased transcriptomic and multiomic profiling of both seminoma and normal human male germ cells, we highlight molecular similarities and differences between seminoma and migratory/pre-migratory PGCs. We uncovered both genetic and epigenetic signatures distinguishing seminoma from normal developmental lineages, which has profound diagnostic and treatment implications for male germline cancer and infertility. Moreover, we developed an in vitro seminoma model using PGCLCs with i(12p), which partly recapitulates activation of several signaling pathways, thus serving as a foundation for construction of a bona fide in vitro seminoma model.

## Methods

### RESOURCE AVAILABILITY

#### Lead contact

Further information and requests for resources and reagents should be addressed to, and will be fulfilled by, the lead contact, Kotaro Sasaki.

#### Materials availability

hiPSCs generated in this study are available from the Lead Contact with a completed Materials Transfer Agreement.

#### Data and code availability

Accession numbers generated in this study are GSE256162 (10x Chromium scRNA-seq data, 10x Chromium Multiome data and bulk RNA-seq data). The codes used for pseudotime analysis are available at GitHub repository: (https://github.com/chengkeren/seminoma).

## Method details

### Collection of human samples

Urogenital organs at human fetuses at 4-16 wpf, many used in previously published stides^17,20^, were obtained from donors undergoing elective abortion at the Fukuzumi Obstetrics and Gynecology Clinic, Hokkaido, Japan, and the Gynecology Clinic, University of Pennsylvania, Philadelphia, PA (Table S1). Embryo ages were determined by the ultrasonographic measurement of crown rump length. Testicular tissues at 35 wpf and 4M were collected within 24 hrs of the demise through autopsy service at the Children’s Hospital of Philadelphia, Philadelphia, PA. Testicular wedge tissues from clinically prepubertal boys 5, 8 and 12 years of age were collected at the Children’s Hospital of Philadelphia. Samples were collected upon informed consent. Experimental procedures were approved by the IRB at the University of Pennsylvania, Children’s Hospital of Philadelphia, and Hokkaido University. Sex of the fetuses was determined by sex-specific polymerase chain reaction (PCR) on genomic DNA isolated from trunk tissues by using primers for the ZFX/ZFY loci^68^ Urogenital and testicular biopsy tissues were dissected in RPMI-1640 medium (Gibco) with the assistance of a dissection microscope. For the 4 wpf embryo, the entire urogenital ridge containing the coelomic epithelium, mesonephros and a portion of the proximal mesentery were isolated^20^. For samples at 5-8 wpf, complete isolation of the gonads from the surrounding structures could not be made in certainty due to small size and partial disruption during the clinical procedure, allowing occasional contamination of non-gonadal tissues (e.g., adrenal tissues). Likewise, one adrenal sample at 6 wpf contained testicular somatic cells/germ cells, which were included in the study.

Seminoma tissues were derived from two patients undergoing orchiectomy surgery at University of Pennsylvania (The Hospital of the University of Pennsylvania and Penn Presbyterian Medical Center). The patients did not undergo chemotherapy prior to orchiectomy. Thin slices of seminoma tissues were procured at the pathology department for histologic analysis, scRNA-seq and/or multiome analysis. The diagnosis of pure seminoma was made by pathologists based on morphologic features and immunophenotypes diagnostic of seminoma (i.e., POU5F1^+^KIT^+^CD30^−^). Human samples used in this study are listed in Table S1.

### Establishment and Feeder free culture of human iPSCs

Dermal fibroblasts obtained from PKS patients were reprogrammed using the CytoTune-iPS 2.0 Sendai Reprogramming Kit (Thermofisher) following manufacturer’s instructions. Briefly, fibroblasts were plated 4 x10^4 in one well of 6 well plate and cultured in mouse embryonic fibroblast [MEF] medium (DMEM containing 10% Fetal bovine serum [FBS]) one day before infection. On the first day (Day 0), three Sendai virus reprogramming vectors containing expression cassettes of human KOS (*KLF4*, *OCT3/4*, *SOX2*), *L-MYC* or *KLF4* were added to culture at a 5:5:3 ratio. Medium was changed to fresh MEF medium at days 1, 3 and 5. At day 7, cells were harvested and re-plated onto 6 well plate coated with iMatrix-511 silk (Nacalai USA) at 1:10 split in MEF medium. At day 8, medium was switched to Stemfit Basic03 (Ajinomoto) containing 10 ng/ml basic FGF. Between day 14 and 28, individual colonies were handpicked and transferred to a new well containing iMatrix 511 silk. These cells were passaged 2-3 times until they were cryopreserved as a master cell bank.

One vial from the master cell bank was thawed and subsequently cultured for additional ∼15 passages on 6-well plates coated with iMatrix-511 Silk in StemFit Basic04CT (complete type) medium at 37 °C under 5% CO_2_. For passaging, iPSCs at day 6–7 after passaging were treated with a 1:1 mixture of TrypLE Select (Life Technologies) and 0.5 mM EDTA/phosphate-buffered saline (PBS) for 12-15 min at 37 °C to dissociate them into single cells. 10 μM Y-27632 (Tocris) was supplemented in culture medium 24 h after passage of iPSCs. These cells were cryopreserved in at least 20 cryovials as working cell bank from which cells used for PGCLC induction experiments or authentication are derived.

### Authentication of cell lines

For counting chromosomes, iPSCs were incubated with 100 ng/ml of KaryoMAX Colcemid solution (Gibco) for 10 h in culture medium. Cells were dissociated using TrypLE Select and incubated with hypotonic solution (75 mM KCl) for 30 min at 37 °C. The cells fixed by Carnoy’s solution were dropped onto glass slides in a moisty chamber to prepare chromosomal spread. The number of chromosomes was counted using DAPI staining, and the cell lines confirmed to be harboring 46 or 47 chromosomes were further analyzed for G-banding using Cell Line Genetics (Madison, WI). We verified that all cell lines banked are mycoplasma free using MycoAlert mycoplasma detection assay (Cambrex). All experiments used passage-number matched iPSCs (between p18 and p25). Under our feeder free culture method, iPSCs can be stably maintained without loss of pluripotency^69^. For PKS iPSC lines, we applied the simplified pluripotency verification method by assessing transcript levels for pluripotency associated genes (i.e., *POU5F1*, *NANOG*) by qPCR, and morphological analyses of colonies, as we reported previously^42^. To verify trilineage differentiation, 1 x 10^6 iPSCs suspended in 100 μl of PBS containing 30% v/v Matrigel (Corning) were injected subcutaneously in NCG mice under the back skin. Teratoma were harvested at 1-2 months after transplantation then processed for histologic analysis upon fixation in 10% neutral buffered formalin.

### Antibodies

The primary antibodies used in this study included mouse anti-TFAP2C (Santa Cruz Biotechnology, sc-12762), mouse anti-POU5F1 (Santa Cruz Biotechnology, sc-5279), mouse anti-SALL4 (Biocare Medical, CM 384A), goat anti-SOX17 (R&D Systems, AF1924), goat anti-DDX4 (R&D Systems, AF2030), rabbit anti-CTAG2 (Novus Biologicals, NBP2-49659), rabbit anti-MAGEC2 (Abcam, ab209667), rabbit anti-MKI67 (Abcam, ab-15580), Brilliant Violet 421-conjugated mouse anti-human CD49f (Biolegend, 313623) and APC-conjugated mouse anti-CD326 (Biolegend, 324207). The secondary antibodies included Alexa Fluor 488 conjugated donkey anti-rabbit IgG (Life Technologies, A21206), Alexa Fluor 488 conjugated donkey anti-mouse IgG (Life Technologies, A32766), Alexa Fluor 568 conjugated donkey anti-mouse IgG (Life Technologies, A10037), Alexa Fluor 568 conjugated donkey anti-rabbit IgG (Life Technologies, A10042), and Alexa Fluor 647 conjugated donkey anti-goat IgG (Life Technologies, A21447).

### IF analyses on paraffin sections

IF analyses for human testicular samples were performed on paraffin sections as previously described^20^. Tissues were placed in 10% neutral buffered formalin within 2 hrs after the surgical recovery. Samples submerged in formalin were incubated ∼24 hrs at room temperature with gentle rocking. After dehydration, tissues were embedded in paraffin, serially sectioned at 4 mm thickness with a microtome (Leica RM2035) and placed on glass slides (Superfrost plus or Platinum Pro). Paraffin sections were then de-paraffinized with xylene. Antigen retrieval was conducted by treatment of sections with HistoVT one (Nacalai USA) for 35 min at 90°C and then for 10 min at room temperature. The slides were washed twice with PBS, then incubated with blocking solution (5% donkey serum, 0.2% Tween 20 and 1X PBS) for 1 hr at room temperature. The primary antibody incubation was performed overnight at 4°C, and slides were washed with PBS six times (20 min each) followed by incubation with secondary antibodies in bocking solution and 1 μg/mL DAPI for 50 min. Slides were subsequently washed 6 times with PBS before being mounted in Vectashield mounting medium (Vector Laboratory) for confocal microscopy analysis (Leica, SP5-FLIM inverted). Confocal images were processed using Leica LasX (version 3.7.2).

### Co-IF and In situ hybridization (ISH) on paraffin sections

ISH on formalin-fixed paraffin-embedded sections was performed using the ViewRNA ISH Tissue Assay Kit (Thermo Fisher Scientific) with gene-specific probe sets for human *BICD1* (VA1-3001814-VT), *ESRG* (VA1-3020729-VT), and *PRDX2* (VA1-3004726). Experiments were conducted according to the manufacturer’s instructions (incubation with the pretreatment buffer for 12 min, the protease treatment for 6 min 30 sec, FastRed as a chromogen)^20^. Slides were subsequently subjected to IF for TFAP2C as described above except antigen retrieval step. Slides were mounted in Vectashield mounting medium for confocal microscopic analysis.

### 10x Genomics single-cell RNA-seq library preparation

Samples used for scRNA-seq are listed in Table S1. Chromium Single Cell 3L Reagent Kit (v3.1 chemistry) was used. For seminoma and testicular samples beyond 15 wpf, tissues were minced by scissors and dissociated into single cells by Multi Tissue Dissociation Kit 1 (Miltenyi Biotec) according to the manufacturer’s protocol. For testicular or other fetal tissues below 12 wpf, isolated tissue fragments were washed twice with PBS followed by being minced with scissors in 500 μl of 0.1% trypsin/EDTA solution, then incubated for 9 min at 37°C with gentle pipetting every 3 min. After quenching of the reaction by addition of 500 μl of dissection medium (10% fetal bovine serum in Dulbecco’s Modified Eagle Medium), cell suspensions were strained through a 70 µm nylon cell strainer to remove cell clumps then centrifuged at 220 g for 5 min. Cell pellets were resuspended in 0.1% BSA in PBS and counted. All samples were stained with trypan blue and confirmed to be over 80% viable. Cells were loaded into Chromium microfluidic chips to generate single cell gelbead emulsions with a Chromium controller (10x Genomics) according to the manufacturer’s protocol. Gelbead emulsion-RT was performed with a C1000 Touch Thermal Cycler equipped with a deep-well head (Bio-Rad). All subsequent cDNA amplification and library construction steps were conducted according to the manufacturer’s instruction. Libraries were sequenced with a NextSeq 500/500 high output kit v2 (150 cycles) (FC-404–2002) on an Illumina NextSeq 550 sequencer.

### Mapping reads of 10x Chromium scRNA-seq and data analysis

Raw data were demultiplexed with the mkfastq command in Cell Ranger (v7.1.0) to generate Fastq files. Trimmed sequence files were mapped to the reference genome for humans (GRCh38) provided by 10X Genomics using Cell Ranger mkfastq. Read counts were obtained by using cellranger count^70^.

Secondary data analyses were performed in Python with Scanpy 1.8.1^71^, or in R (v.4.1.0) with the Seurat (v.4.3.0), SeuratObject (v4.1.3), SeuratWrappers (v0.3.2), tidyverse (v.1.3.2), destiny (v3.10.0), pheatmap (v1.0.12), plot3D (v1.4) and Matrix (v.1.5-1) packages and Excel (Microsoft). UMI count tables were first loaded into R by using the Read10x function in Seurat, and Seurat objects were built from each sample. Cells with less than 200 genes, an aberrant high gene count above 7000, or total mitochondrial genes above 15% were filtered out. Of the ∼89477 cells for which transcriptomes were available, 72257 cells passed quality-control dataset filters and used for downstream analysis. We detected ∼3000 median genes/cell at a mean sequencing depth of ∼62400 k reads/cell (Fig. S1A). Samples were combined using Seurat merge function. The effects of mitochondrial genes, cell cycle genes, were regressed out by SCTransform during normalization in Seurat. Batch effect was removed using Harmony. Mitochondrial genes and cell cycles genes were excluded during Cell Clustering, dimension reduction and trajectory analyses. Cells were clustered by a shared nearest neighbor (SNN) modularity optimization based clustering algorithm in Seurat. Clusters were annotated on the basis of previously characterized marker gene expression by using the FeaturePlot function and the gene expression matrix file. Then, annotated cell clusters were used for downstream analyses. Dimensional reductions were performed with top 3000 highly variable genes and first 20 principal components by using Seurat. Differentially expressed genes (DEGs) in different clusters were calculated by MAST or area under curve (AUC) integrated in Seurat findallmarkers function, with a threshold of avgLog2FC > 0.25, *p*-value < 0.01. Developmental trajectories of cells were simulated with first 30 PCs and 30 diffusion components by DiffusionMap in destiny and scanpy^72^. Trajectory principal lines were fitted by DiffusionMap. Pseudotime was calculated by dpt in destiny. For LogNormalize Expression (LogExp) in Seurat, genes for each cell were divided by the total counts for that cell and multiplied by the cal factor (10,000), then natural-log transformed using log1p. For Pseudobulk differential expression analysis of scRNA-seq data, DEGs between two groups in scatterplot were identified by edgeR 3.34.1 with applying quasi-likelihood approach (QLF) and the fraction of detected genes per cell as covariate^73^. The DEGs were defined as the genes with FDR < 0.05, *p*-value < 0.05, and Log2-fold change above 1. Expression value was normalized by edgeR using log2(CPM+1) (CPM = Counts Per Million). Cell Cycle was analyzed using CellCycleScoring in Seurat. Data was visualized by ggplot2 and pheatmap. Genes in heatmap were hierarchically clustered by measuring Euclidean distance, scaled by row, then visualized using pheatmap. Gene ontology enrichment was analyzed by DAVID v6.8.

### Library preparation for 10x Chromium single cell multiome ATAC + Gene Expression

Human fetal testicular tissues were minced by scissors and dissociated into single cells by Multi Tissue Dissociation Kit 1 (Miltenyi Biotec) according to the manufacturer’s protocol. Single cell suspensions were resuspended in 0.04% BSA in PBS and then proceed to nuclei isolation according to the low cell input method in the manufacturer’s protocol (10× Genomics), using ice cold lysis buffers consisting of nuclease-free water with final concentrations after dilution: 10mM Tris-HCl (pH7.4, Sigma-Aldrich), 10mM NaCl (Sigma-Aldrich), 3mM MgCl_2_ (Sigma-Aldrich), 0.01% Tween-20 (Thermo Fisher Scientific), 0.01% Nonidet P40 substitute (Sigma-Aldrich), 0.001% digitonin (Thermo Fisher Scientific), 1% BSA (Miltenyi Biotech), 1mM DL-Dithiothreitol (DTT, Sigma-Aldrich), 1U/μl sigma protector RNase inhibitor (Sigma-Aldrich). Diluted nuclei suspensions were mixed with Trypan Blue to determine optimal lysis and nuclei concentration by counting. Finally, nuclei were suspended in nuclei buffer (10X) with 1mM DTT and 1U/μl RNAse inhibitor and then immediately loaded into Chromium Next GEM Chip J with the Chromium Next GEM Single Cell Multiome Reagent Kit to generate GEMs using the Chromium Controller (10× Genomics). GEM incubation was performed in a C1000 Touch Thermal Cycler with 96-Deep Well Reaction Module (Bio-Rad). All subsequent post-GEM cleanup, pre-amplification, ATAC library construction, cDNA amplification and GEX library construction steps were performed according to the manufacturer’s protocol. Libraries were sequenced using a 2 × 150 paired-end sequencing protocol on an NovaSeq 6000 instrument.

### Mapping reads and data analysis for Single Cell Multiome ATAC + Gene Expression

Raw sequencing data of single cell multiome seq was converted to FastQ format using Cellranger. For each sequenced multimodal snRNA-seq library, we performed read alignment to GRCh38 v.3.1.0 (human) reference genome, provided by 10X Genomics, quantification and initial quality control using the Cell Ranger ARC Software (v.1.0.1, 10X Genomics) using default parameters. Cell Ranger filtered count matrices were used for downstream analysis. The downstream analysis was performed using ArchR (v1.0.2)^74^. Generally, cells with Log10(unique fragments) > 3, peak_region_fragments > 3000, peak_region_fragments < 100000, pct_reads_in_peaks > 40, blacklist_ratio < 0.025, nucleosome_signal < 4, TSS.enrichment > 6 were used for downstream analysis. Clustering and the remaining analysis were following default parameters(resolution = 0.2, sampleCells = 10000, n.start = 10, varFeatures = 2500). TSS was extended with upstream 2000bp and downstream 200bp. Genebody was extended with upstream 2000bp and downstream 1000bp. Iterative Latent Semantic Indexing (LSI) was used for dimensional reduction. motif enrichment was performed by HOMER (v4.6). Genome coverage was generated and visualized by ArchR (v3.5.1).

To reduce dimensions of the dataset, a binary datum of accessibility was derived by transforming counts greater than 0 to 1 for remaining peaks. Binarized accessibility was used as input for TF-IDF weighting, using term frequency and smoothed inverse document frequency as weighting scheme. Iterative Latent Semantic Indexing (LSI) was used for dimensional reduction, UMAP and t-SNE were used to project cells into two dimensions.

Peaks were called using MACS2 with default parameters^75^, which is integrated in ArchR. Briefly, cells in each annotated cluster were grouped together and peaks were called by MACS2. Peaks per cell < 500 and > 150000 are excluded. Valid peaks were presented in at least 25 cells, mitochondria chromosome and chromosome Y were excluded duing peak calling. Blacklists annotated by encode project were excluded. Promoter region was defined as 2000bp upstream and 100 downstream of transcription start site. Then, peaks were annotated with ChIPpeakAnno (v3.28.1) using GRCh38 as reference genome. Peaks in distal intergenic regions were subseted by bedtools and used for GREAT GO analysis^76^.

To discover transcription factor dynamics and variation in their motif accessibility we conducted analysis using chromVar^77^. Briefly, we used position weight matrices (PWMs) for 579 known TFs from JASPAR. Transcription factor motif to peak assignments were used in conjunction with counts from 500□bp size fixed cluster-specific peaks to calculate an accessibility deviation *Z*-score for each transcription factor motif/cell pair.

### Integrated analysis of gene expression, gene score and motif enrichment

Multi-group comparisons in gene expression, gene score and motif enrichment analyses were performed using ArchR. Then, genes showing the same up-regulated pattern in three layers of data were selected for generation of integrated heatmap. Gene expression level was normalized by LogNormalize Expression (LogExp) in Seurat, genes for each cell were divided by the total counts for that cell and multiplied by the cal factor(10,000), then natural-log transformed using log1p. The Gene Score model in ArchR was used to generate the gene accessibility score. Briefly, blacklist is excluded during analysis^78^. Gene scores are calculated as follows: tiles within the gene window of a certain gene are identified, and the ones that overlap with another gene region are excluded. Of the remaining tiles, the distance to the gene is calculated and an exponential weighing function is applied to also take into consideration distal regulatory elements. To address the bias resulting from differences in gene size, ArchR applies a separate weight for the inverse of the gene size (1 / gene size) and scales this inverse weight linearly from 1 to a user-defined hard maximum (default of 5). Gene scores can be calculated directly during arrow file creation or can be added later. A gene score matrix was generated for downstream analysis. The getMarkerFeatures and getMarkers function in ArchR (testMethod□=□“wilcoxon”, cutOff□=□“FDR□<□=□0.05”) was used to identify the marker regions/genes for each cluster. Gene-score imputation was implemented with addImputeWeights for data visualization. Ingenuity Pathway Analysis (IPA) was performed by QIAGEN Digital Insight solutions.

### Single cell copy number analysis

Initial CNVs (CNV) were estimated by sorting the analyzed genes by their chromosomal location and applying a moving average to the relative expression values, with a sliding window of 100 genes within each chromosome, as described^79^. To avoid considerable impact of any particular gene on the moving average, we limited the relative expression values to [–3,3] by replacing all values above 3 by a ceiling of 3, and replacing values below –3 by a floor of –3. This was performed only in the context of CNV estimation. This initial analysis is based on the average expression of genes in each cell relative to the other cells and therefore does not have a proper reference to define the baseline. We thus defined the gene expression clusters annotated as PGCs.E, PGCs.L, M.pSpg, Int.pSpg, and T1.pSpg by gene expression as nonmalignant cells, and used the average CNV estimate at each gene across those cells as the baseline.

The final CNV estimate of cell i at position j was defined as:

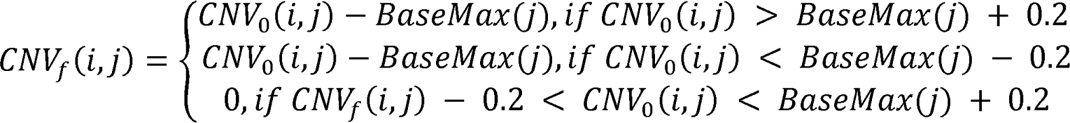

The 6 states correspond to the following CNV events:

- State 1; 0x, complete loss.
- State 2; 0.5x, loss of one copy.
- State 3; 1x, neutral
- State 4; 1.5x, addition of one copy
- State 5; 2x, addition of two copies
- State 6; 3x, essentially a placeholder for >2x copies but modeled as 3x.

### Profiling of TE expression levels from 10X chromium scRNA-seq data

To quantify the expression level of TEs at locus resolution, we first prepared a transcript annotation file (i.e., GTF file) including human TEs, annotated by RepeatMasker and Dfam^80,81^. Reads were mapped and counted using Cell Ranger with the above annotation file. TE loci detected in ≥ 0.5% of the cells were used in the downstream analyses. The read abundances of total TEs, respective TE families, and TE subfamilies were calculated by summing the read counts of TE loci. The expression levels of TEs were normalized as raw counts for each cell divided by the total counts for that cell and multiplied by the scale.factor = 10000. Then natural-log transformed using log1p by Seurat LogNormalize. Dimension reduction analysis of the TE expression profile obtained from scRNA-seq data were performed according to the Seurat flamework (https://satijalab.org/seurat/). The 100 most variably expressed TE subfamilies were selected using the FindVariableFeatures command. After the data were scaled and whitened, dimension reduction analysis was performed using the Uniform Manifold Approximation and Projection (UMAP) and t-Distributed Stochastic Neighbor Embedding (t-SNE). The first ten principal components were used in the analysis. For pseudobulk pair-wise comparison, expression of TEs were normalized by log2(CPM+1), and DEGs were calculated by edgeR, with a definition of log2fold change > 1, p-value < 0.05 and FDR < 0.05. Violin plots and scatterplots were visualized by ggplot2. Heatmaps were generated by pheatmap.

### Quantitative RT-PCR

Quantitative RT-PCR (qPCR) was performed as described previously^42,82^. Briefly, cell pellet were lysed for RNA isolation using RNeasy Micro Kit (Qiagen). cDNA was synthesized using 1 ng of purified RNA and amplified by PCR as described previously ^42,83^. ERCC spike-in RNAs developed by the External RNA Controls Consortium (ERCC; life Technologies) were spiked in the sampled to gauge the transcript copy number per cell. Target cDNAs were quantified using primers (Table S9) and PowerUp SYBR Green Master Mix (Applied Biosystems) with StepOnePlus (Thermo Fisher Scientific). The Log2 scale gene expression values were calculated using ΔCt method normalized to the averaged Ct values of housekeeping genes, *PPIA* and *ARBP*.

### Bulk RNA-seq library preparation

Total RNA of iPSCs and ITGA6^+^EPCAM^+^ PGCLCs were extracted with RNeasy Plus Micro Kit (#74034, QIAGEN). RNA-seq libraries were made using SMRT-Seq HT plus kit (#R400748, Takara) according to the manufacturer’s protocol. Library preparation was performed at the Center for Host-Microbial Interactions at the University of Pennsylvania School of Veterinary Medicine. Briefly, total RNA was quantified using Qubit Fluorometric quantification, and RNA integrity was verified with a TapeStation (Agilent). RNA Integrity Number equivalent (RINe) above 8 was used for downstream libraries preparation. cDNA library was prepared using Clontech SMART-Seq HT Plus Kit (PN R400749) according to the manufacturer’s instructions. cDNA was purified with AMPure XP beads. 75-base pair reads were sequenced on an Illumina NextSeq 500 machine according to the manufacturer’s protocol.

### Bulk RNA-seq data analysis

Raw fastq files were demultiplexed with bcl2fastq2 (v.2.20.0.422). Barcodes and adapters were removed with Trimmomatic (v.0.32). Fastq files were mapped to UCSC human reference genome (GRH38) using STAR (2.7.10a). The raw gene count table was generated with featurecounts, and weakly expressed genes were filtered with edgeR with the filterByExpr function with default parameters. Briefly, the raw counts were normalized to library size, and then genes with counts per million above 10 were included in downstream analysis. DEGs were analyzed with edgeR (v.3.36.0) with log2fold change >1, p-value <0.05, and FDR <0.05. Reads per kilobase per million (RPKM) values were calculated in edgeR, and the gene length was obtained from the UCSC table browser. Downstream data analyses and visualization were performed with R (v.4.1.0). Hierarchical clustering was performed with hclust in R (v.4.1.0).

## ACKNOWLEDGMENTS

We thank Drs. Leslie King and Jerome F. Strauss III for carefully reviewing the manuscript and providing insightful comments. We thank Drs. Walter J. Storkus, Jumpei Ito, Louisa C. Pyle, Katherine L. Nathanson, Bangjin Kim, and Kosuke Izumi and members of Sasaki lab for the discussion of this study. We acknowledge the Comparative Pathology Core at the University of Pennsylvania School of Veterinary Medicine for the preparation of paraffin sections.

This work was supported in part by Open Philanthropy projects (2019-197906, 10080664, 10095457) to K.Sasaki.

## AUTHOR CONTRIBUTIONS

K. S conceived the project, designed the experiments, and wrote the manuscript. K.C. and A.R. edited the manuscript. Y.S. and R.Y. generated and analyzed PGCLCs. Y.S.H., I.D.K., J.P.G., P.L., X.L., P.M.P., R.L.L., T.F.K., S.R. assisted in human sample procurement. K.C. prepared samples for scRNA-seq. K.C. and E.W. contributed to the analyses of scRNA-seq and multi-omics.

## DECLARATION OF INTERESTS

The authors declare no competing interests.

---

**Figure S1. Single cell RNA-seq profiles of human male germline development, related to Figure 1.**

**(A)** Violin plots showing quality control metrics used in scRNA-seq analyses. Samples are colored as in (B).

**(B)** UMAP plot showing all cells from human testes and a urogenital ridge used to isolate germ cells for trajectory analysis in (E). Each dot represents a single cell, colored according to sample origin as in Table S1.

**(C)** UMAP plot as in (B) projected with cell types identified based on Seurat clustering analysis and annotated on the basis of marker gene expression and DEGs as in (D)^20^.

**(D)** Heatmap showing the averaged expression of DEGs identified from a multi-group comparison among cell types defined in (C) (log fold change > 0.25, p < 0.01). Log-normalized expression value is scaled by row with Z-score transformation. Key DEGs are listed on the right.

**(E)** UMAP plot showing subsetting and re-clustering of germ cells defined in (C).18 cell types are defined based on Seurat clustering analysis and annotated on the basis of marker genes and DEGs as in (F).

**(F)** Heatmap showing the averaged expression of DEGs identified by a multi-group comparison among subtypes of germ cells as in (E) (log fold change > 0.25, p < 0.01). Log-normalized expression value is scaled by row with Z-score transformation. Representative marker genes are listed on the right.

**Figure S2. Gene expression dynamics during human male germline development, related to Figure 1.**

**(A-I)** Scatter plots comparing the averaged gene expression between indicated cell types as defined in Fig. 1A. Differentially expressed genes (DEGs) (absolute log2 fold change above 1, p-value < 0.05 and FDR < 0.05) are highlighted in colors and the number of DEGs are shown. Representative genes and their GO enrichments for DEGs are shown beneath each plot.

**Figure S3. Transcriptome landscape of seminoma, related to Figure 2.**

**(A)** H&E staining of a seminoma tissue (Se1). Scale bar, 50μm.

**(B)** IF of seminoma sections (Se1) for SALL4, SOX17 (green), NANOG, TFAP2C (red) counterstained with DAPI (white). Merged images are shown on the right.

**(C)** Quality control metrics of scRNA-seq for two seminoma samples (Se1 and Se2). Mt, mitochondrial genes; ribo, ribosomal genes; hb, hemoglobin genes.

**(D, E)** UMAP plot showing of all cells based on computationally aggregated scRNA-seq data obtained from two seminoma samples. Each dot represent a single cell, colored according to cell cycle scoring (D) or sample origin (E).

**(F)** Scatter plot comparing averaged gene expression of all cells between two seminoma samples. Pearson Correlation coefficient is shown in red.

**(G)** Expression of key marker genes projected on t-Distributed Stochastic Neighbor Embedding (t-SNE) embedding. Each cell type is outlined by dotted lines.

**(H)** Heatmap showing the averaged expression of DEGs identified by a multi-group comparison among lymphocyte cell types as in Fig. 2G (log fold change > 0.25, p < 0.01).

Representative genes and their GO enrichments are shown on the right.

**(I, J)** t-SNE embedding showing subtypes of dendritic cells and macrophages. Macrophages and DCs in Fig.2C are subsetted and re-clustered with new annotation assigned based on marker gene expression (J). ICMs, inflammatory cytokine-enriched macrophages; LAMs, lipid-associated macrophages; cDC1s, classical dendritic cell type 1; cDC2s, classical dendritic cell type 2; A/M-cDCs, activated/mature classical dendritic cells; pDCs, plasmacytoid dendritic cells.

**(K)** Heatmap showing the averaged expression of DEGs identified by a multi-group comparison among indicated macrophage and DC subtypes as defined in (I) (log-fold change > 0.25, p < 0.01). Log-normalized expression value is scaled by row with Z-score transformation. Representative markers genes and enriched gene ontology terms were listed on the right.

**(L)** Top 5 DEGs among cell types defined in **(I).** Dot size represents the percentage of cells expressing the gene. The color of dots denote the averaged expression of the gene.

**Figure S4. Transcriptomic comparison of seminoma and normal human male germ cells, related to Figure 3.**

**(A)** Expression of seminoma-specific (*ESRG*, *LIM2*), pluripotency-associated (*POU5F1*) and T1.pSpg/Spg (*PIWIL4*) markers projected on DiffusionMap embedding as in Fig. 3A.

**(B)** Pearson correlation of transcriptomes among PGCs.E, PGCs.L, M.pSpg and seminoma.

**(C)** Venn diagram showing genes shared among iPSC-derived male germ cells and seminoma cells. For iPSC-derived male germ cells, upregulated DEGs of each cell type over other in vitro cell types are used, which are obtained by multi-group comparison of scRNA-seq data^17^. For seminoma cells, upregulated DEGs of seminoma compared to other TME cells are used. DEGs with average log2-fold change >0.25, p <0.01, FDR <0.01 are used. Set sizes (total number of DEGs used) and intersection sizes (genes overlapped with indicated cell types) are shown as bar graph. Note that upregulated genes in seminoma show the largest overlap with those in PGCLCs.

(**D, E**) Scatter plot comparing averaged expression between seminoma and PGCs.L (D) or between seminoma and M.pSpg (E). DEGs (absolute log2 fold change > 1, p-value < 0.05 and FDR < 0.05) are highlighted in colors and the number of DEGs are shown. Representative genes and their GO enrichments are shown beneath each plot.

**(F)** Violin plots showing representative upregulated genes shared between seminoma and various developmental stages of human male germ cells.

**(G)** Venn diagrams showing overlap of seminoma markers (upregulated DEGs of seminoma vs TME cells) with markers of indicated human male germ cells in vivo (upregulated DEGs of each cell type vs others obtained through multi-group comparison).

**(H)** Co-ISH/IF of two seminoma samples (Se1, Se2) for TFAP2C (green), *BICD1*, *ESRG* (red) counterstained with DAPI (white). Merged images are shown on the right. Bar, 50 μm.

**(I)** Heatmap showing DEGs obtained by multigroup comparisons of PGCs.E, PGCs.L, M.pSpg and seminoma. Enriched GO terms and representative genes are listed on the right. DEGs were defined as average log2 fold change > 0.25, p-value < 0.01. Heatmap shows the Z-score normalized gene expression by row.

**Figure S5. Intratumor heterogeneity of seminoma, related to Figure 3.**

**(A, B)** UMAP plot of all seminoma cells subsetted from Fig.2C and re-clustered. Cells are colored according to 5 clusters identified by Seurat clustering analysis (A) or by sample origin (B).

**(C)** Single-cell reconstruction of developmental trajectories of seminoma using DiffusionMap, related to Fig. 3A. For comparison, PGCs.E, PGCs.L and M.pSpg are included in the analysis.

**(D)** Marker genes for each subtype of seminoma projected on UMAP plot as in (A). Pan-seminoma markers and other key negative markers are also shown on the left.

**(E)** Violin plots showing representative genes with progressive increase along the trajectory in (C) and Fig. 3A, B.

**(F)** Heatmap showing averaged expression of DEGs identified from a multi-group comparison among subtypes of seminoma (log fold change > 0.25, p < 0.01). Representative genes and enriched GO terms are listed on the right.

**(G)** Ingenuity signaling pathway analysis of seminoma subtypes and PGCs.E. Dot size represents −log10(p-value), color denotes Z-score of pathway activation/repression.

**Figure S6. Simultaneous profiling of single cell ATAC-seq and single cell RNA-seq of human male testis in second trimester fetus, related to Figure 4.**

**(A, B)** Per-cell quality control. Fragment size distribution plot in ATAC-seq showing enrichment around 100 and 200 bp, representing nucleosome-free and mono-nucleosome-bound fragments (A). Plot showing transcription start site (TSS) enrichment scores and unique nuclear fragment numbers (B).

**(C)** Violin plots showing indicated quality control metrics.

**(D)** UMAP plot showing all cells of a second trimester human gonad analyzed based on nuclear RNA expression (scRNA-seq) (top left), chromatin accessibility (scATAC-seq) (bottom left) and both metrics combined (LSI combined) (right). Cells are colored according to cell types annotated by marker genes, DEGs and DGASs as in (H) and (I).

**(E)** Heatmap showing differential accessible regions among each cell type.

**(F)** Heatmap showing Pearson correlation of open chromatin status among different cell types.

**(G)** Genome browser screenshot of *NANOG* and *TCL1A* loci in indicated cell types.

Peak-to-gene linkage shows strength of correlation between chromatin accessibility and gene expression.

**(H)** Heatmap showing DEGs obtained from multi-group comparison of cell types. Key DEGs are shown on the right.

**(I)** Heatmap showing DGASs obtained from multi-group comparison of cell types. Key markers are shown on the right.

**(J)** Heatmap showing Pearson correlation of gene activity scores and gene expression in different cell types.

**(K)** Integrated analysis of gene expression (RNA), gene activity scores and motif enrichment (Motif). Genes whose values in all of the above categories are significantly higher in one cell cluster over the other clusters are shown as “active” transcription factors.

**Figure S7. Simultaneous profiling of single cell ATAC-seq and single cell RNA-seq in M.pSpg and T1.pSpg, related to Figure 4.**

**(A)** Germ cells in a second trimester human testis identified in Fig. S6D were subsetted and re-clustered by Seurat clustering analysis according to nuclear RNA expression (scRNA-seq), chromatin accessibility (scATAC-seq) or both metrics combined (LSI-combined). Cells are colored based on cell types annotated as M.pSpg (red) and T1.pSpg (blue) based on marker gene expression as in (C).

**(B)** Pseudotime projected on the UMAP as in (A).

**(C)** Projection of gene expression or gene activity scores of indicated markers of PGCs/M.pSpg (top) or T1.pSpg (bottom) onto UMAP as in (B). *ASB9*, a marker of Int.pSpg is shown in the middle.

**(D)** Dynamics of gene expression, gene activity scores and motif enrichment along pseudotime defined in (B). Genes showing coordinated patterns between gene expression and Motif enrichment (left) or between gene activity scores and motif enrichment (right) are shown.

**(E)** HOMER motif enrichment analysis using peaks defined in (F), and the average expression of genes binding to motifs. Dot size denotes the enrichment significance, which was normalized as −log10(p-value). Gene expression was log normalized. Blue color indicates no mRNA expression.

**(F)** Venn diagram showing open chromatin regions unique to M.pSpg or T1.pSpg or overlapping between two.

**(G)** Stacked barplots showing genomic distribution of open chromatin regions identified in (F).

**(H)** Genome browser screenshots of *TFAP2C* and *NANOG* loci.

**Figure S8. Integrated multi-omics profiling of seminoma cells and their tumor microenvironment, related to Figure 4**.

**(A)** UMAP plot showing all cells in seminoma tissues clustered according to nuclear RNA expression (scRNA-seq), chromatin accessibility (scATAC-seq) or both metrics combined (LSI-combined). Clusters are annotated based on marker gene expression and DEGs as in (C, D).

**(B)** Pearson correlation between Gene activity scores and gene expression in each cell type.

**(C)** LSI-UMAP projections of multi-layer features of seminoma specific TFs. Gene (RNA) expression, gene activity scores and corresponding motif enrichment of indicated transcription factors are shown on the same UMAP embedding.

(**D, E**) K-means clustered heatmap showing average gene expression (D) or gene activity scores (E) of genes showing differential values among cell types. Each row represents a gene and each column represents a cell type. Selected genes and enriched GO terms in each list are shown on the right.

**(F)** Integrated analyses of gene expression, gene activity scores, and motif enrichment.

Genes whose values in all of above categories are significantly higher in one cell type over the others are shown as “active” transcription factors.

**(G)** Genome browser screenshots of *TFAP2C* and *NANOG* loci in seminoma cells and their tumor microenvironment cells.

**Figure S9. Isochromosome 12p in seminoma and establishment of iPSCs from individuals with Pallister-Killian syndrome, related to Figure 6**.

**(A)** Fluorescent in situ hybridization (FISH) of a seminoma sample (Se2) using *PKP2* (targeting chromosome 12p11, red) and D12Z3 (targeting centromere of chromosome 12, green) as probes. Arrows indicate the presence of isochromosome 12p.

**(B)** Copy number variations (CNVs) analysis of seminoma. Normal germ cells (PGCs.E, PGCs.L, M.pSpg, Int.pSpg and T1.pSpg) were used as reference, sex chromosomes were removed during analysis. Dark red defines gain of copy number, and deep blue defines loss of copy number. Chromosome 12p is labeled and representative genes in this genomic region are listed below. Key marker genes expressed in seminoma cells are highlighted in red.

**(C)** Demographics of PKS patients and established iPSC lines. Mosaicism was assessed by FISH on dermal fibroblasts.

**(D)** Phase contrast images and karyotypes of iPSCs derived from PKS patients.

**(E)** Growth rate of iPSCs bearing i(12p) and their isogenic control (46XY) established from the same PKS patient (PKS19).

**(F)** qPCR quantification of pluripotency-associated and house-keeping genes of isogenic iPSCs with/without i(12p). Expression values of spike-in RNAs (*ERCC1806*, *ERCC452*, *ERCC56*) are shown as controls. qPCR was performed using amplified cDNA derived from 1ng RNA as described in Methods.

**(G)** H&E images of a representative teratoma derived from 19-1iPSCs bearing i(12p). using +i(12p) hiPSCs. Scale bar, 100 μm.

## SUPPLEMENTARY DATASETS, see separate Excel documents

**Supplementary Table S1.** Sample information used in this study.

**Supplementary Table S2.** DEGs among all cell types in human samples, related to Fig. S1

**Supplementary Table S3.** DEGs among germ cell types in vivo related to Fig. S1

**Supplementary Table S4.** Number of cells per germ cell type in vivo across sample stages, related to Fig. 1

**Supplementary Table S5.** DEGs of all cell types in seminoma tissues

**Supplementary Table S6.** DEGs of lymphocyte subtypes in seminoma TME.

**Supplementary Table S7.** DEGs of macrophages and dendritic cell subtypes in seminoma TME.

**Supplementary Table S8.** Differentially expressed TEs identified from a pairwise comparison between seminoma and TME cells, related to Fig. 5

**Supplementary Table S9.** Primers used in this study

## Notes

### Competing Interest Statement

The authors have declared no competing interest.

