## Supplementary figures and images for "Defining the cellular origin of seminoma by transcriptional and epigenetic mapping to the normal human germline"

### Fig. S1

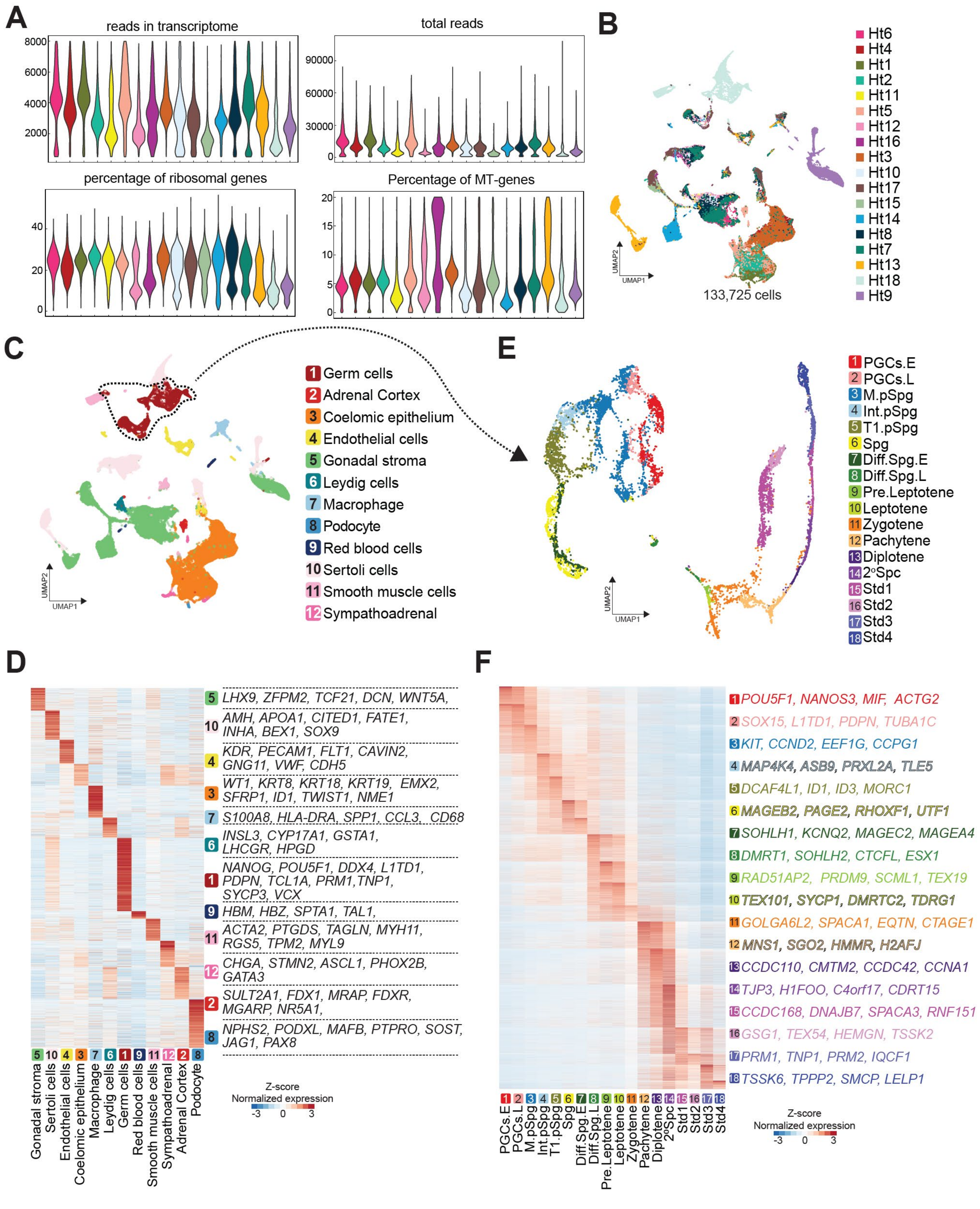

### Fig. S3

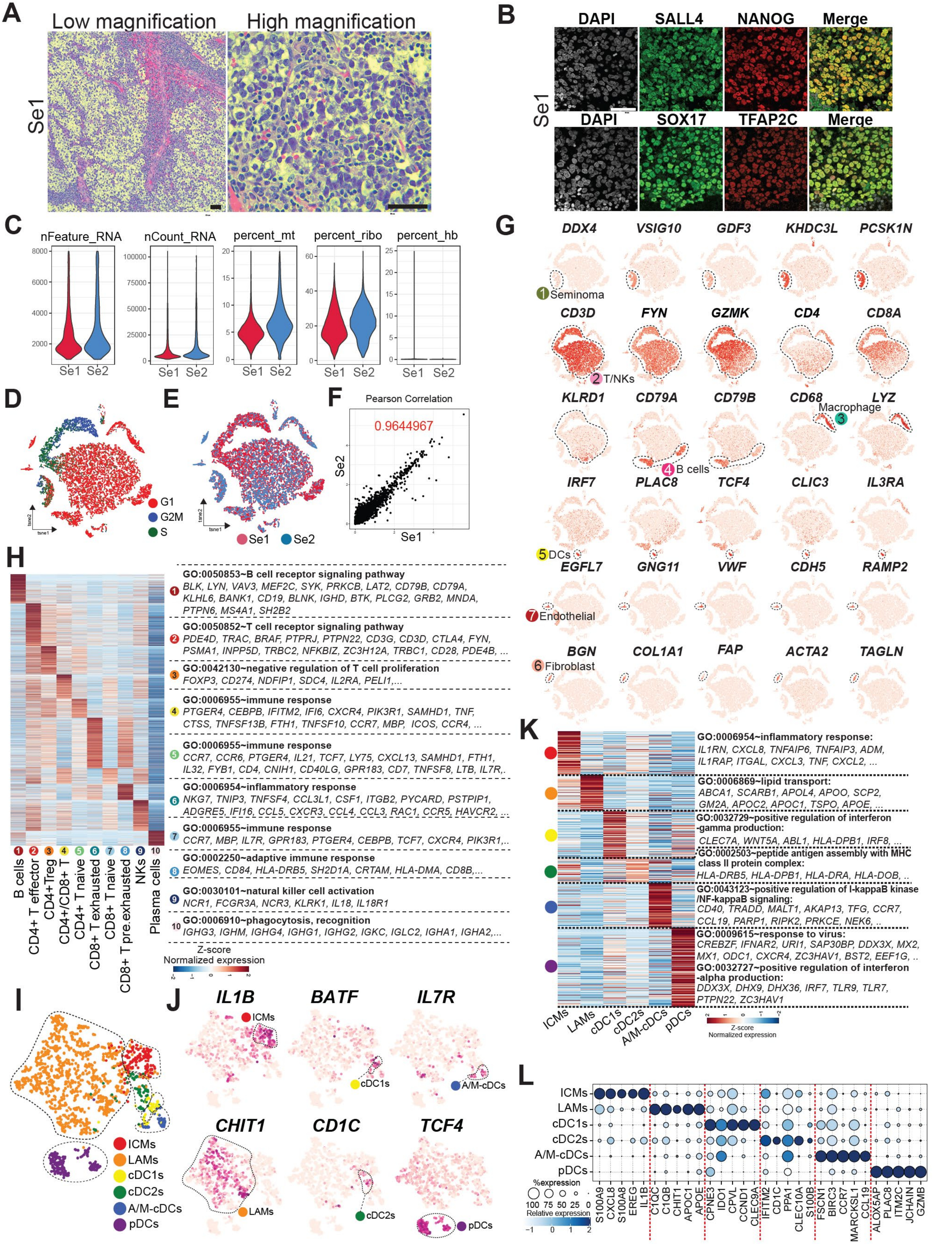

### Fig. S4

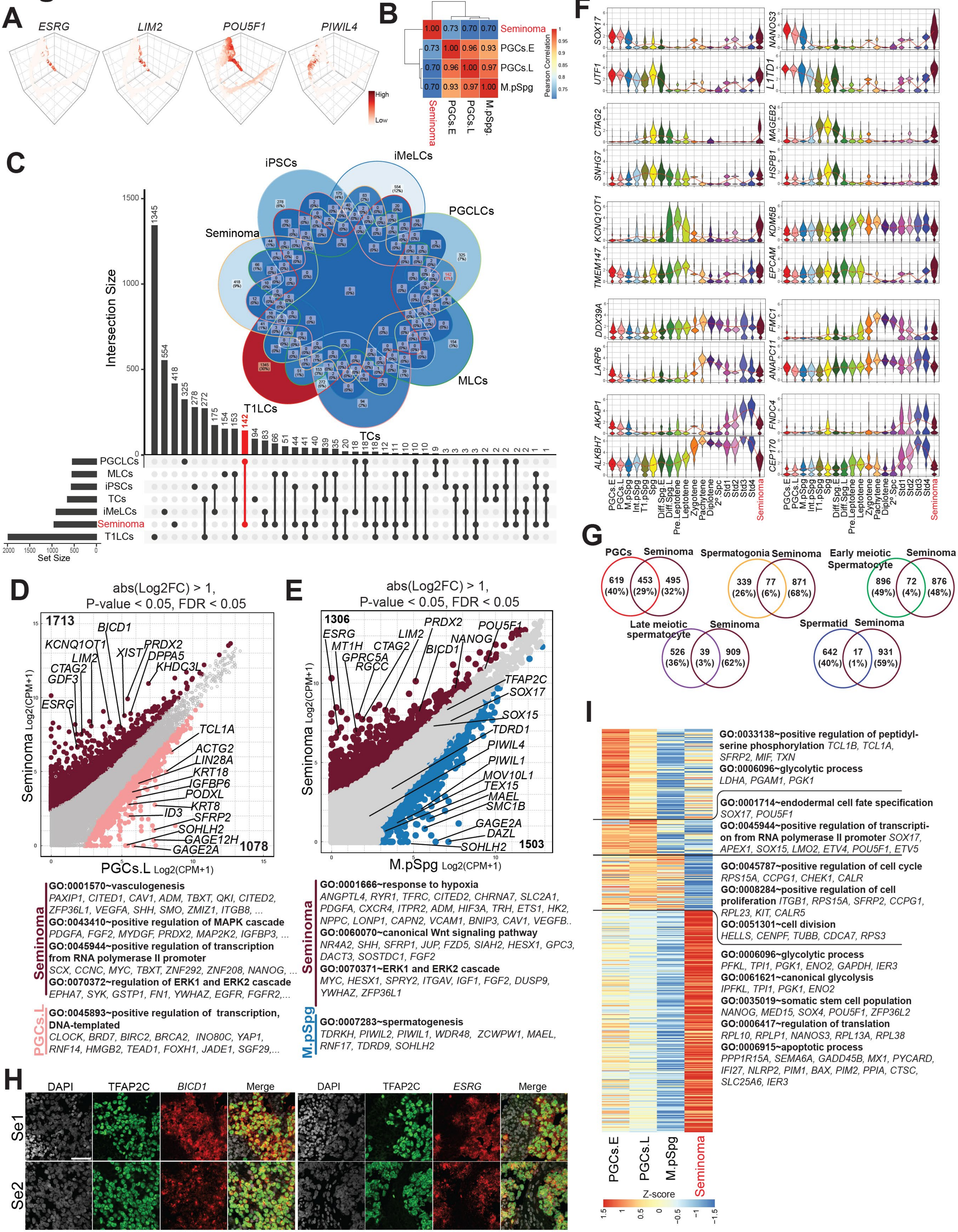

### Fig. S5

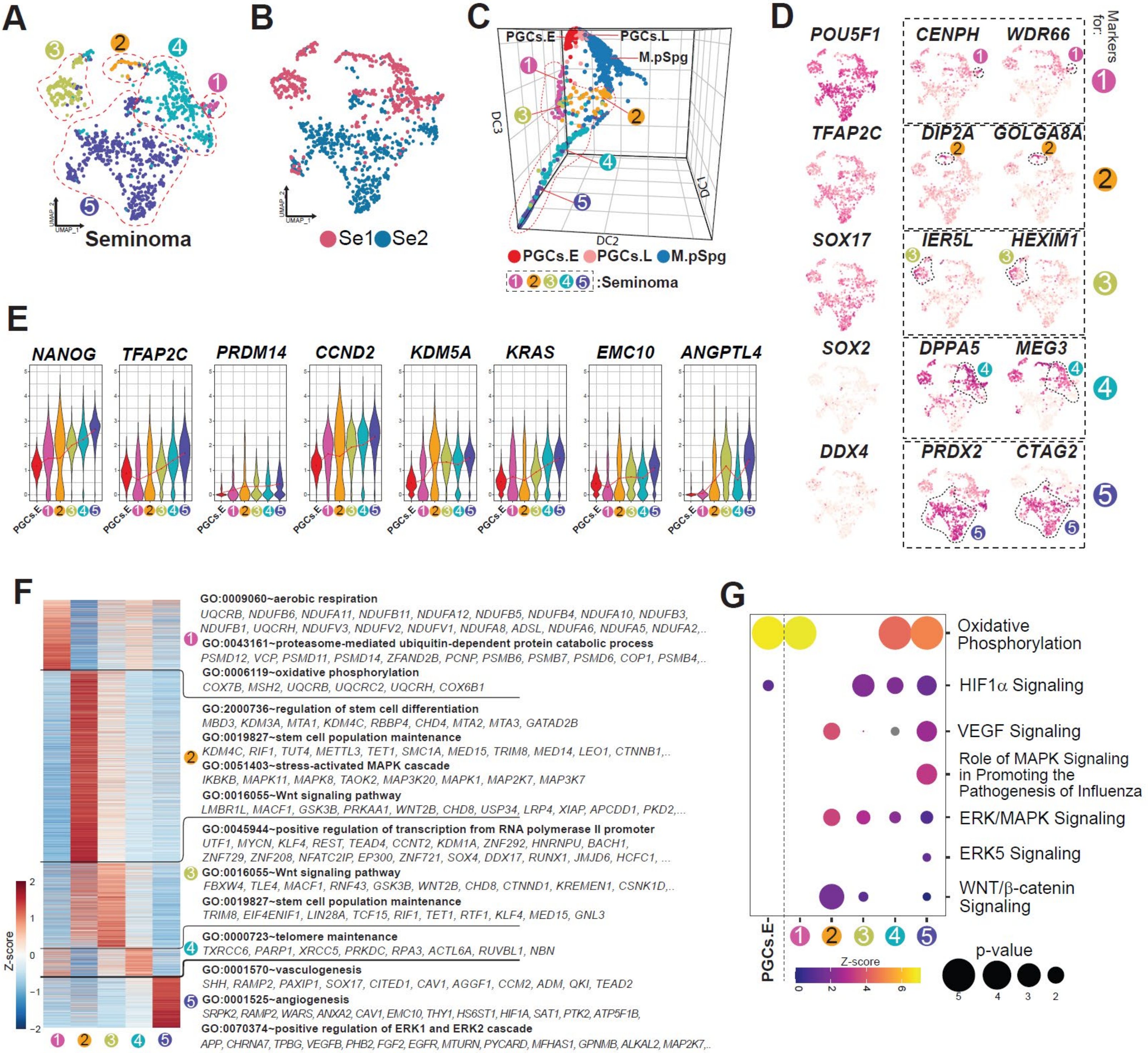

### Fig. S6

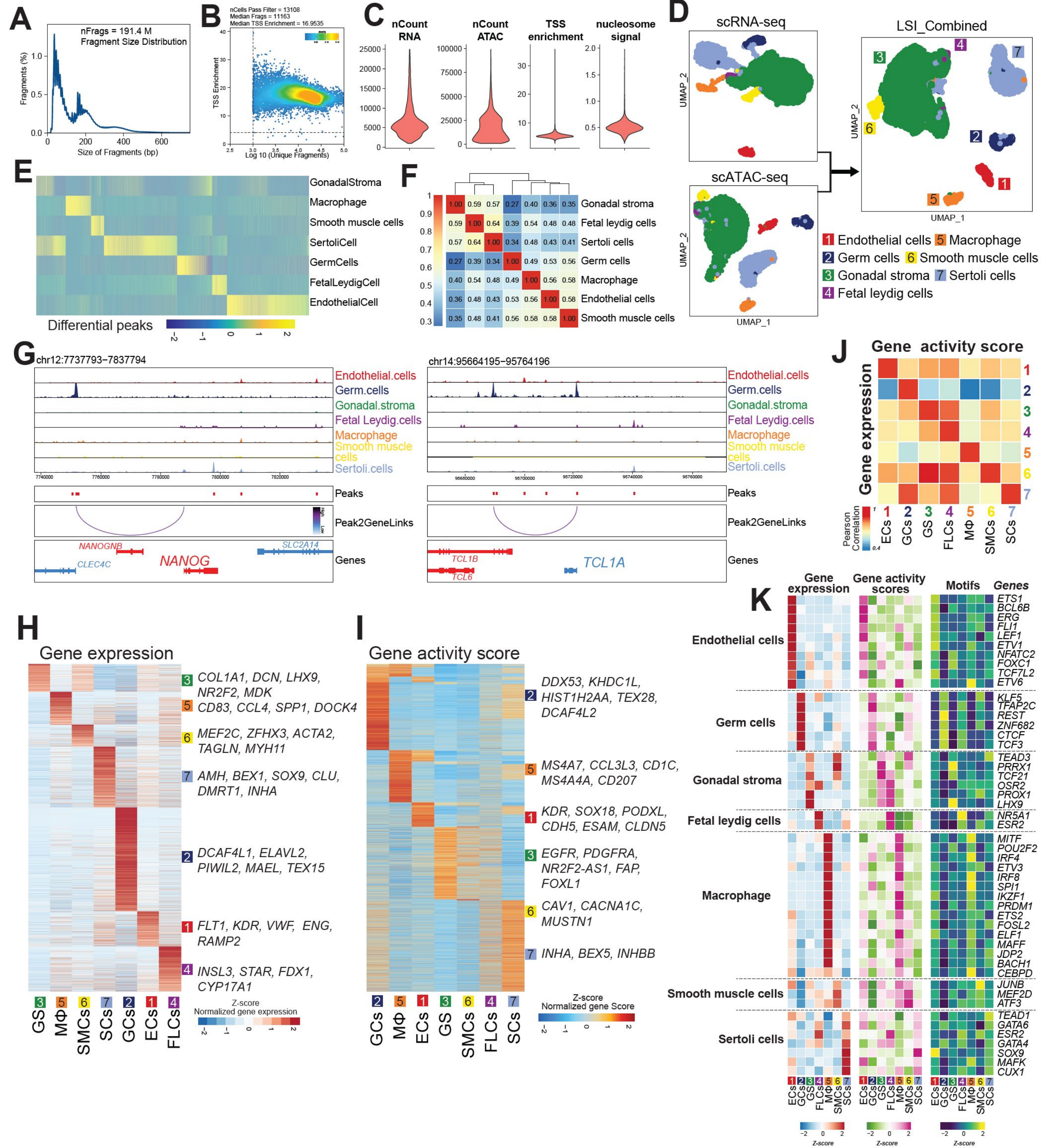

### Fig. S7

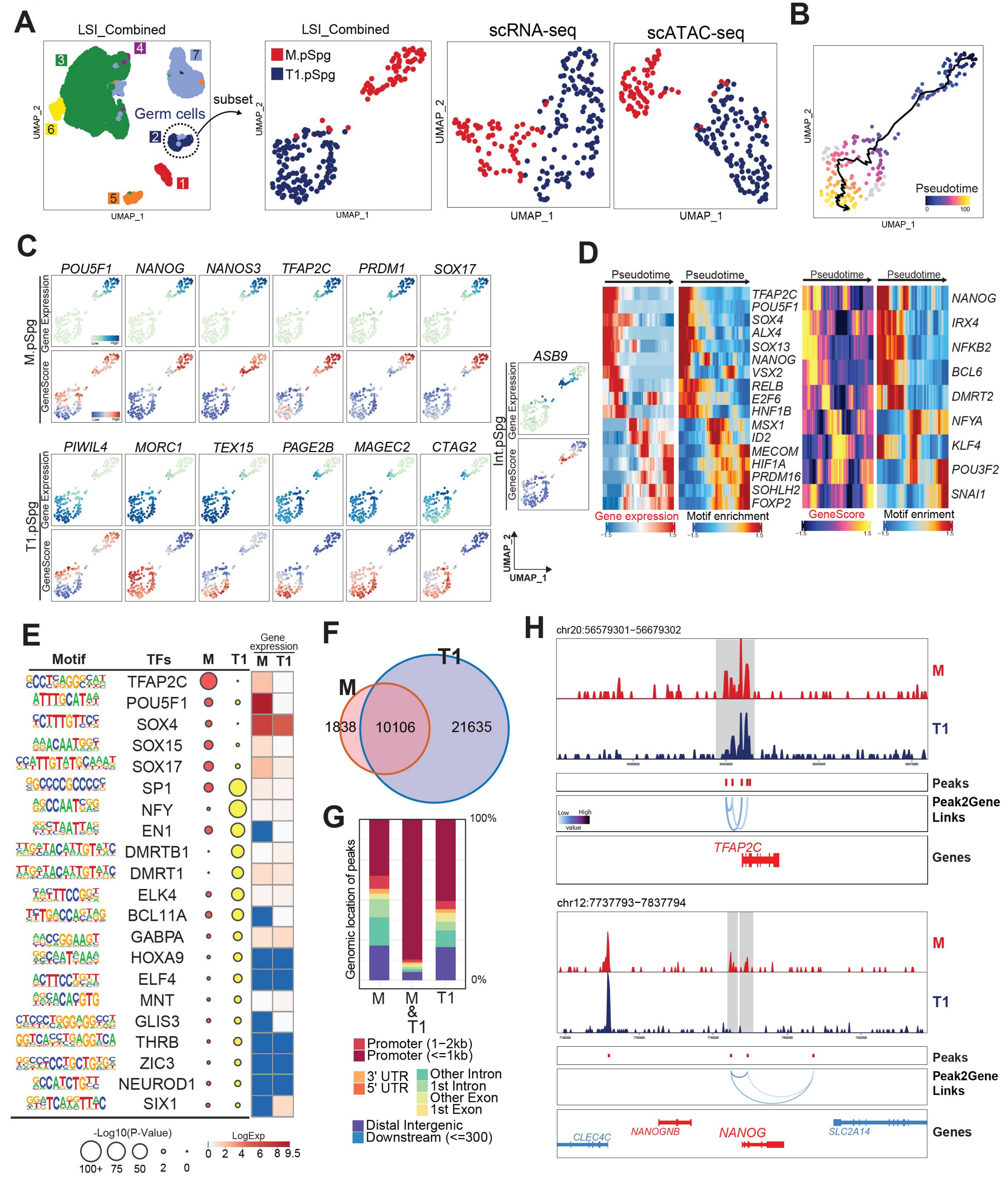

### Fig. S8

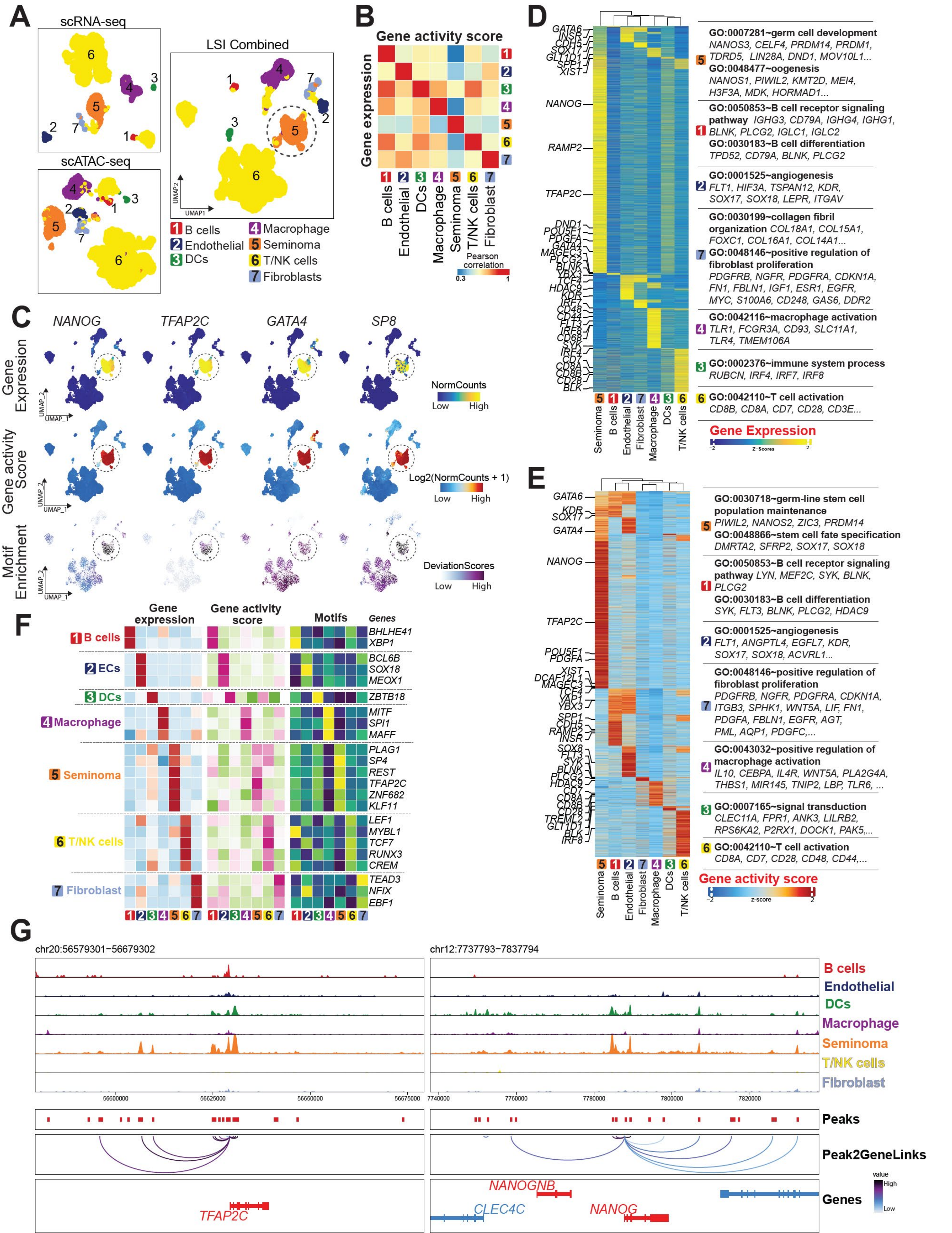
