## Supplementary material for "Defining the cellular origin of seminoma by transcriptional and epigenetic mapping to the normal human germline": Fig. S2

### Figure S2

A

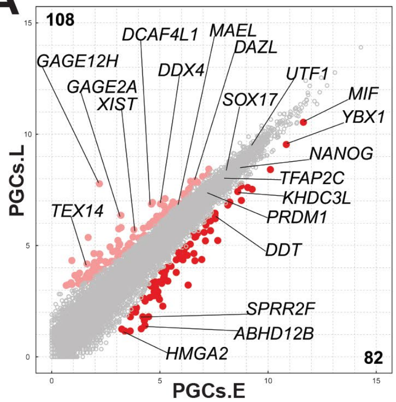


GO:0030334~regulation of cell migration
*BEX4, LAMA4, NEXN*
GO:2000036~regulation of stem cell population maintenance *HMGA2, PTN*

GO:0007140~male meiosis
*DDX4, MAEL, TDRD9, TEX14*
GO:0007283~spermatogenesis
*DAZL, ZCWPW1, DDX4, MAEL, TDRD9, RNF17, CLOCK...*

B

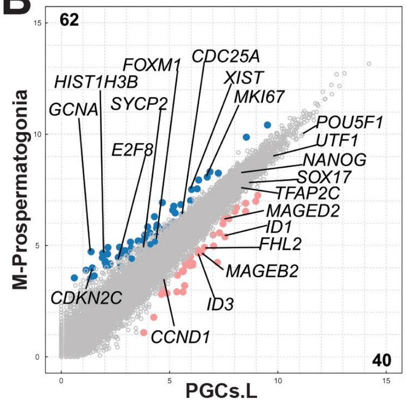


**GO:0000122~negative regulation of transcription from RNA polymerase II promoter** *IRX1, EID1, CCND1, MAGED1, ID1, MAGEB2, ID3, MAGED2, MAGEB1*

**GO:0007140~male meiosis**
*TEX14, TEX15*
**GO:0007049~cell cycle**
*MKI67, CDC25A, SPC25*

C

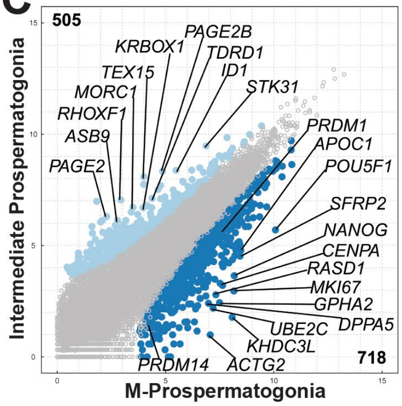


**GO:0051301~cell division**
*ERCC6L, CCNF, KIF14, BUB1B,TUBA1C, TUBA1B, CCND2, CCNA2, UBE2C, CENPF, WEE1,...*
**GO:0007281~germ cell development**
*DAZL, LIN28A, MSH2, NANOS3, BAX,...*
**GO:0043046~DNA methylation involved in gamete generation**
*PIWIL2, PIWIL4, DDX4, TDRD5, CTCFL, TDRD1,...*
**GO:0034587~piRNA metabolic process**
*ASZ1, MAEL, MOV10L1, TDRD6, PLD6, TDRD5, ...*

D

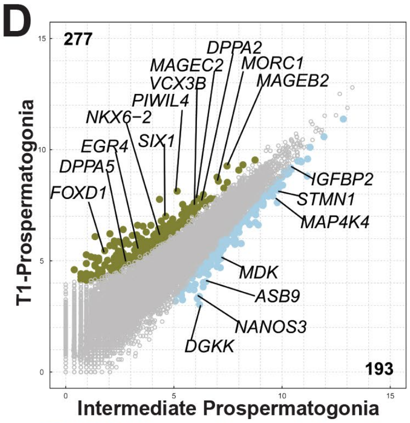


**GO:0017148~negative regulation of translation** *NANOS2, NANOS3, SHMT1, CALR, GAPDH*
**GO:0032981~mitochondrial respiratory chain complex I assembly**
*NDUFS7, NDUFA11, NDUFB11, NDUFS5, ...*

**GO:0034587~piRNA metabolic process**
*TDRD1, PIWIL1, PIWIL4, ASZ1, TDRD6, ...*
**GO:0045944~positive regulation of transcription from RNA polymerase II promoter** *SIX1, GATA2, MSX1, FOXD1, EGR4, HOXA10, HES5, PAX6, POU3F2..*

E

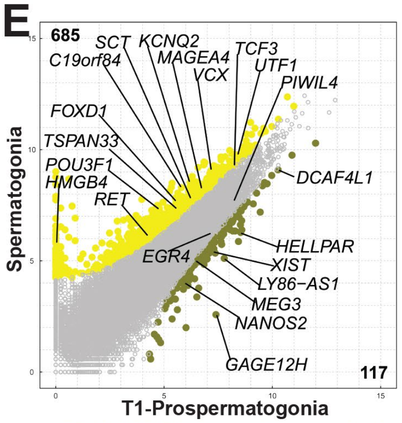


**GO:0006325~chromatin organization**
*L3MBTL1, SETD5, KMT2A, CHD9, PWWP3A, ZNF518A, TET1, CHD2*
**GO:0006306~DNA methylation**
*ATF7IP, KMT2A, ATRX*

**GO:0000165~MAPK cascade**
*RET, MAP2K2, IGFBP4, HSF1, NRTN, RBX1, MAPK3...*
**GO:0009060~aerobic respiration**
*NDUFB7, NDUFB6, NDUFB10, ...*

F

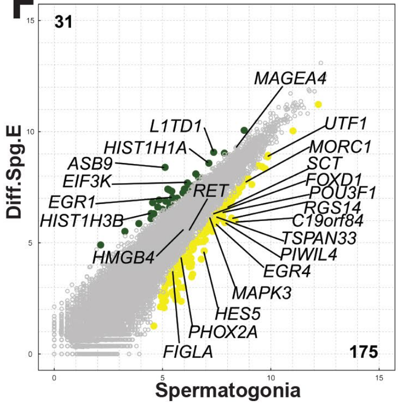


**GO:0045944~positive regulation of transcription from RNA polymerase II promoter** *ZNF296, FOXD1, EGR4, FIGLA, UTF1, PHOX2A, HES5, MAPK3...*
**GO:0031047~gene silencing by RNA**
*PIWIL4, SRRT, MORC1*

**GO:0045944~positive regulation of transcription from RNA polymerase II promoter**
*EGR1, TSTD1, STOX1...*

G

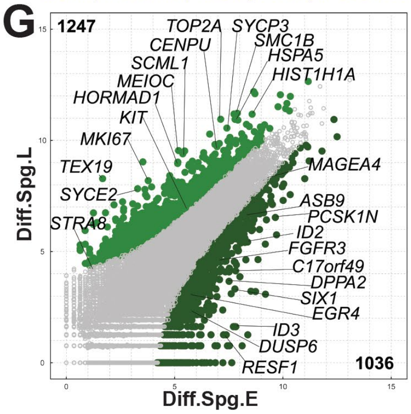


**GO:0030509~BMP signaling pathway**
*DLX5, FST, SMAD9, SMAD6, BMP6, SMAD7, FKBP8, HES1, ROR2, ...*
**GO:0000165~MAPK cascade**
*RET, DUSP5, MAP2K2, IGFBP4,DUSP6, MAPK12, KRAS, FGFR3*
**GO:0006281~DNA repair**
*MDC1, BRCA1, BRCA2, SMC3, UFL1, CHEK1, KMT5B, TOPBP1, TEX15, ...*
**GO:0035234~ectopic germ cell programmed cell death**
*SYCE3, SYCP2, PRKDC, KIT, TEX11, CASP2*

H

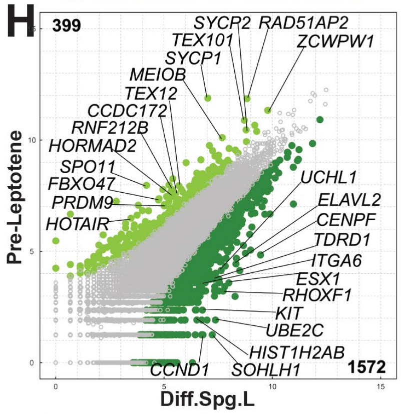


**GO:0006281~DNA repair**
*UHRF1, PNKP, DOT1L, NUDT1, XPC, TTF2, PARK7, UNG, MBD4, SUPT16H,...*
**GO:0008354~germ cell migration**
*ITGB1, TGFB1, DMRT1, KIT, TGFBR1*

**GO:0007129~synapsis**
*MAJIN, PRDM9, SYCP1, DMC1, ZCWPW1, RNF212B, MAEL, MEIOB...*
**GO:0007141~male meiosis I**
*SPO11, DMC1, DMRTC2, MEIOB, SLC25A31, HSPA2, MYBL1*

I

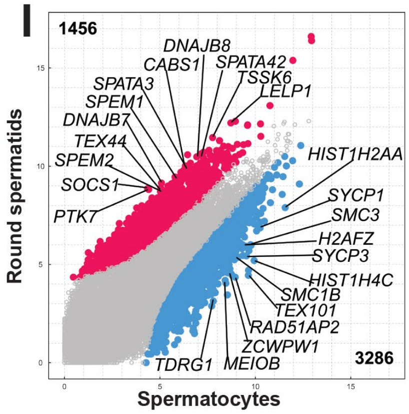


**GO:0007140~male meiosis**
*TDRKH, DDX4, SYCP2, TDRD9, FANCA, TEX14, HSPA2, MLH3, TEX19, TDRD12, MEIOB,...*
**GO:0051321~meiotic cell cycle**
*PIWIL2, PIWIL1, YTHDC2, SMC3, AURKC, BOLL, HORMAD2, HORMAD1, FAM9B, ...*

**GO:0030317~flagellated sperm motility**
*SMCP, TEX43, ARMC12, TEKT5, SPEM1, CABS1, PRM3*
**GO:0007342~fusion of sperm to egg plasma membrane**
*C16ORF92, IZUMO1, SPATA46*
