## Supplementary material for "Defining the cellular origin of seminoma by transcriptional and epigenetic mapping to the normal human germline": Fig. S9

### Figure S9

A

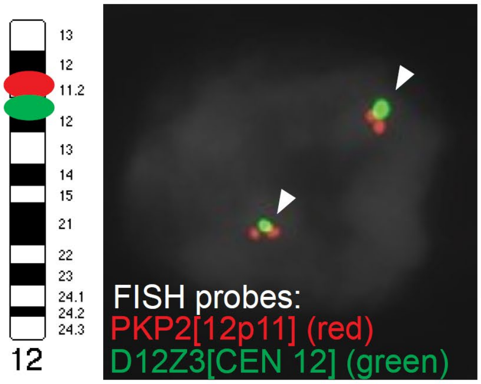


B

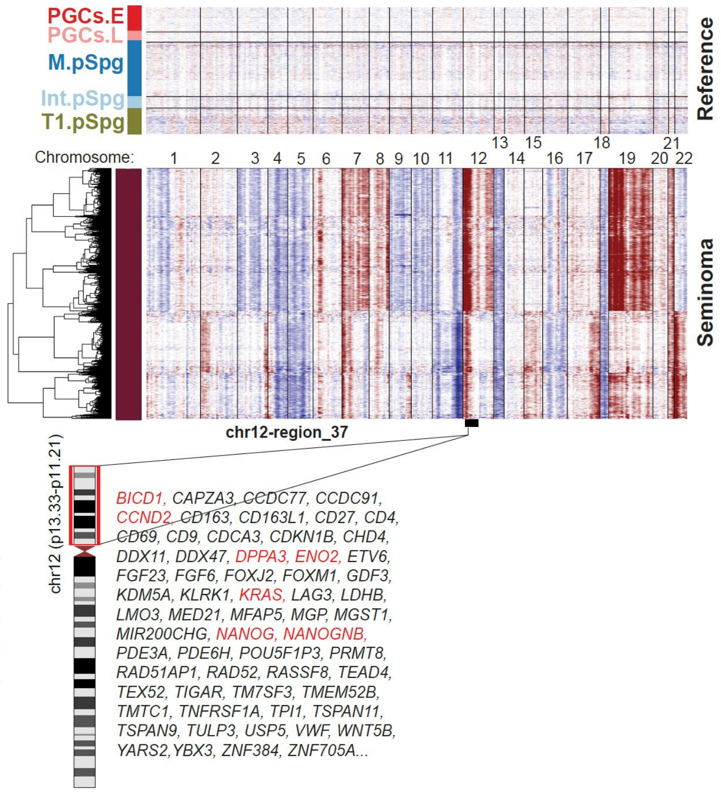


C

| PKS # | Age | % mosaicism | Sex | 46 XY iPSCs | 47 XY, +i(12p) |
|---|---|---|---|---|---|
| PKS12 | 5 weeks | 65% | Male | 12-6 | 12-1, 12-7 |
| PKS19 | 6 months | 90% | Male | 19-3 | 19-1 |
| PKS44 | 9 months | 96% | Male | None | 44-4 |

D

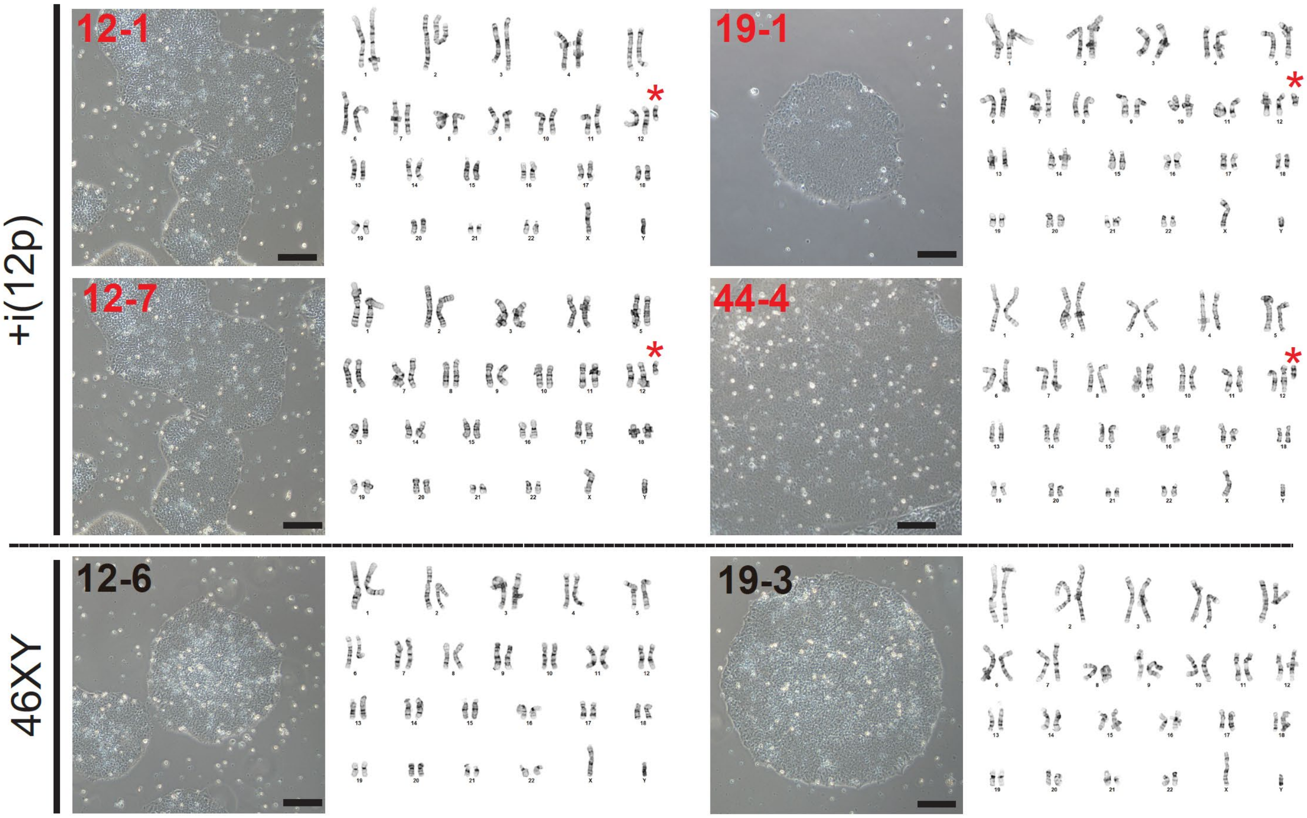


E

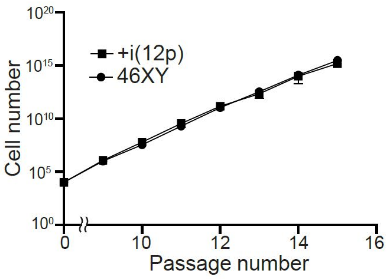


F

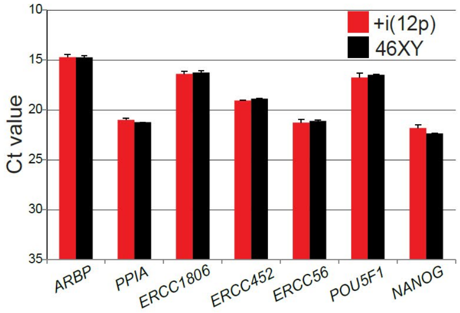


G

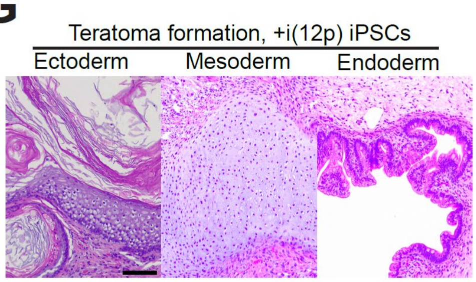
